# Genetic regulation and targeted reversal of lysosomal dysfunction and inflammatory sterol metabolism in pulmonary arterial hypertension

**DOI:** 10.1101/2024.02.26.582142

**Authors:** Lloyd D. Harvey, Mona Alotaibi, Hee-Jung J. Kim, Yi-Yin Tai, Ying Tang, Wei Sun, Wadih El Khoury, Chen-Shan C. Woodcock, Yassmin Al Aaraj, Claudette M. St. Croix, Donna B. Stolz, Ji Young Lee, Mary Hongying Cheng, Tae-Hwi Schwantes-An, Ankit A. Desai, Michael W. Pauciulo, William C. Nichols, Amy Webb, Robert Lafyatis, Mehdi Nouraie, Haodi Wu, Jeffrey G. McDonald, Caroline Chauvet, Susan Cheng, Ivet Bahar, Thomas Bertero, Raymond L. Benza, Mohit Jain, Stephen Y. Chan

## Abstract

Vascular inflammation critically regulates endothelial cell (EC) pathophenotypes, particularly in pulmonary arterial hypertension (PAH). Dysregulation of lysosomal activity and cholesterol metabolism have known inflammatory roles in disease, but their relevance to PAH is unclear. In human pulmonary arterial ECs and in PAH, we found that inflammatory cytokine induction of the nuclear receptor coactivator 7 (*NCOA7*) both preserved lysosomal acidification and served as a homeostatic brake to constrain EC immunoactivation. Conversely, NCOA7 deficiency promoted lysosomal dysfunction and proinflammatory oxysterol/bile acid generation that, in turn, contributed to EC pathophenotypes. *In vivo*, mice deficient for *Ncoa7* or exposed to the inflammatory bile acid 7α-hydroxy-3-oxo-4-cholestenoic acid (7HOCA) displayed worsened PAH. Emphasizing this mechanism in human PAH, an unbiased, metabolome-wide association study (N=2,756) identified a plasma signature of the same NCOA7-dependent oxysterols/bile acids associated with PAH mortality (*P*<1.1x10^-6^). Supporting a genetic predisposition to NCOA7 deficiency, in genome-edited, stem cell-derived ECs, the common variant intronic SNP rs11154337 in *NCOA7* regulated NCOA7 expression, lysosomal activity, oxysterol/bile acid production, and EC immunoactivation. Correspondingly, SNP rs11154337 was associated with PAH severity via six-minute walk distance and mortality in discovery (N=93, *P*=0.0250; HR=0.44, 95% CI [0.21-0.90]) and validation (N=630, *P*=2x10^-4^; HR=0.49, 95% CI [0.34-0.71]) cohorts. Finally, utilizing computational modeling of small molecule binding to NCOA7, we predicted and synthesized a novel activator of NCOA7 that prevented EC immunoactivation and reversed indices of rodent PAH. In summary, we have established a genetic and metabolic paradigm and a novel therapeutic agent that links lysosomal biology as well as oxysterol and bile acid processes to EC inflammation and PAH pathobiology. This paradigm carries broad implications for diagnostic and therapeutic development in PAH and in other conditions dependent upon acquired and innate immune regulation of vascular disease.

**One Sentence Summary:** Pulmonary arterial hypertension pathophenotypes arise from allele-specific NCOA7 regulation of lysosome function and inflammatory oxysterol generation, as demonstrated by genomic and metabolomic association studies coupled with genetic and pharmacologic mechanistic evidence.

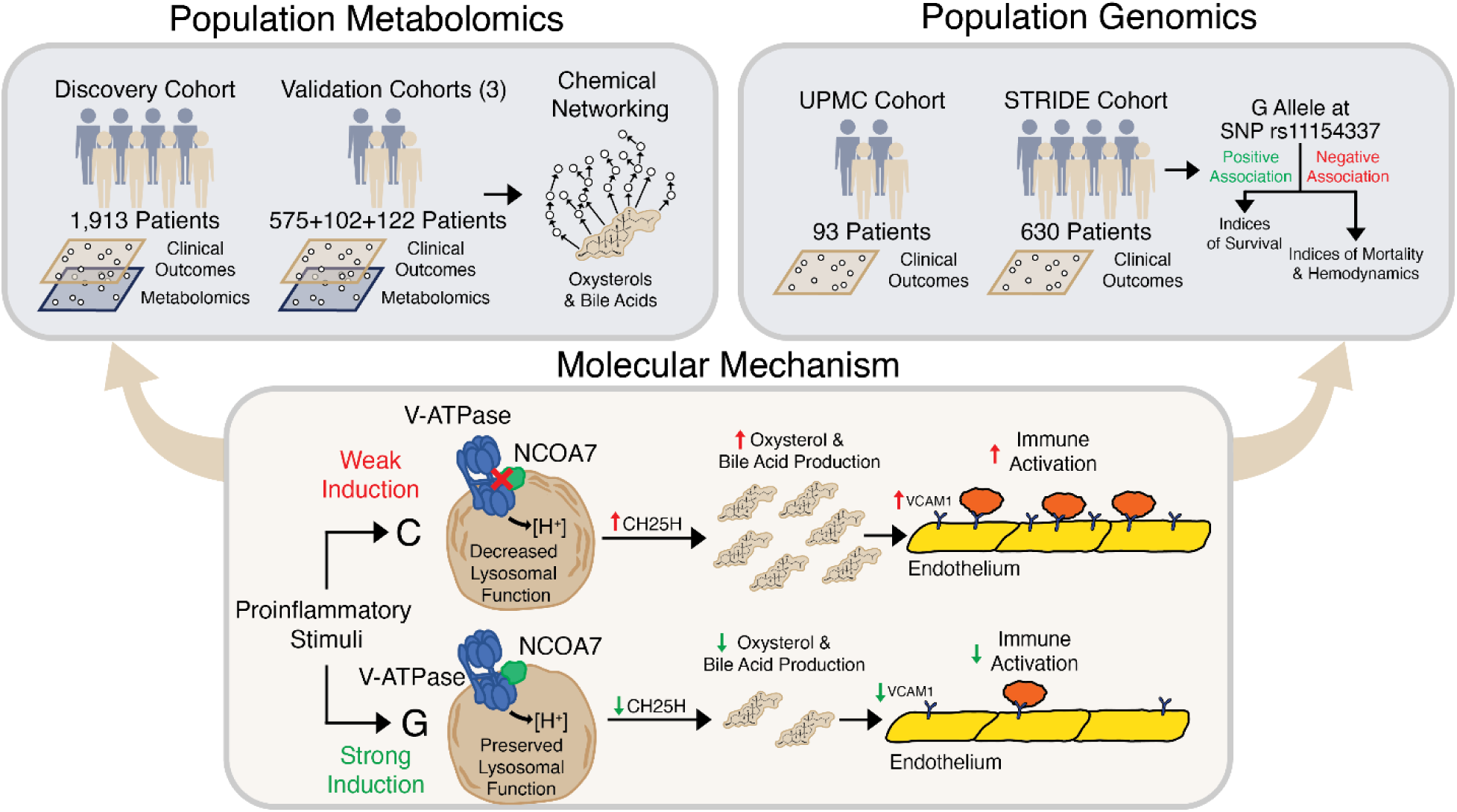
Graphical abstract.

## Introduction

Vascular inflammation critically regulates endothelial cell (EC) integrity and function across multiple vascular disease states including atherosclerosis (*1*), hypertension (*2*), stroke (*3*), and sepsis (*4*). Within the lung, inflammation of the endothelium is a prominent feature of acute lung injury (*5*), pathogen-mediated processes such as SARS-CoV-2 infection (*6*), and pulmonary arterial hypertension (PAH). Of pulmonary vascular disorders, PAH represents among the most morbid and mortal of conditions—a deadly, enigmatic, and chronically progressive disease characterized by complex vessel remodeling and poorly defined molecular origins (*7*). In the absence of definitive knowledge regarding mechanisms driving PAH pathobiology, substantial attention and debate has centered on the possibility that EC inflammation plays a causative role in PAH rather than simply serving as a marker of disease severity or overall risk (*8*).

Although molecular accelerators and brakes are known to regulate responses to inflammation as part of homeostatic maintenance of cellular function (*9*), the specific levers that control EC inflammation are incompletely described. Lysosomal activity is increasingly appreciated as a principal regulator of inflammation (*10*), and dysfunctional lysosomal activity has been observed particularly in the context of PAH (*11*). Central to the maintenance of lysosomal enzyme function is proper acidification of the luminal space, which is mediated by the vacuolar H^+^ ATPase (V-ATPase) family (*12*), and loss of this hydrolytic capacity leads to lysosomal storage disorders that have reported in pulmonary vascular phenotypes (*13–16*). The nuclear receptor co-activator 7 (*NCOA7*) directly binds and modulates V-ATPase activity (*17–19*) to control endolysosomal function, which has documented function in controlling bacterial and viral pathogen entry (*17, 20, 21*), renal tubular acidification (*22*), and neuronal function (*18*). NCOA7 is upregulated in human ECs by proinflammatory stimuli (*23*) and in PAH lung tissue (*24*), but a causative mechanism connecting NCOA7 to cardiopulmonary vascular disease has not been defined.

Several lines of evidence have suggested that lysosomal dysfunction carries relevance to PAH. Downstream of acidification, lysosomes carry pH-sensitive, hydrolytic enzymes responsible for the breakdown of cellular waste and macromolecular trafficking (*25*). The lysosomal-mediated breakdown of cellular waste is connected to autophagy—a process that may be relevant in PAH (*26*). Notably, loss of lysosomal hydrolase activity leads to the accumulation of oxysterols and bile acids (*27*), which are bioactive molecules upregulated in the plasma and lungs of PAH patients (*28–30*). Oxysterols and bile acids influence cholesterol biosynthesis and cell membrane properties, driving critical cellular defenses in adaptive and innate immunity (*31*). At the level of the endothelium, these molecules are capable of EC immunoactivation (*32*) that contributes to peripheral vascular diseases, such as atherosclerosis and hypertension (*33*). Intriguingly, as reported in our parallel study (*34*), an unbiased plasma metabolomic analysis of 2,756 PAH patients identified a metabolome-wide association (adjusted *P*<1.1x10^-6^) of glucuronidated oxysterols and downstream bile acids with clinical PAH disease severity and mortality. As such, given the inherent connection of NCOA7 to both lysosomal biology and a metabolome-wide association of lysosome-derived oxysterol and bile acid levels with PAH disease outcomes, we sought to determine whether NCOA7 controls oxysterol and bile acid metabolism, inflammatory pulmonary EC pathophenotypes, and the development of PAH—thus offering a mechanistic explanation for the observed metabolomic association with PAH severity (*34*).

## Results

An overview of the study design is provided in **Fig. 1A**. The first stage of investigations involved multi-level interrogation of NCOA7 regulation of function, followed by studies of NCOA7 in relation to EC immunoactivation and PAH pathophenotypes. Genetic regulation of NCOA7 was then studied in relation to cellular traits and clinical outcomes. Finally, a pharmacologic NCOA7 activating agent was designed, synthesized, and shown to achieve reversal of PAH phenotypes.

**Fig. 1.**
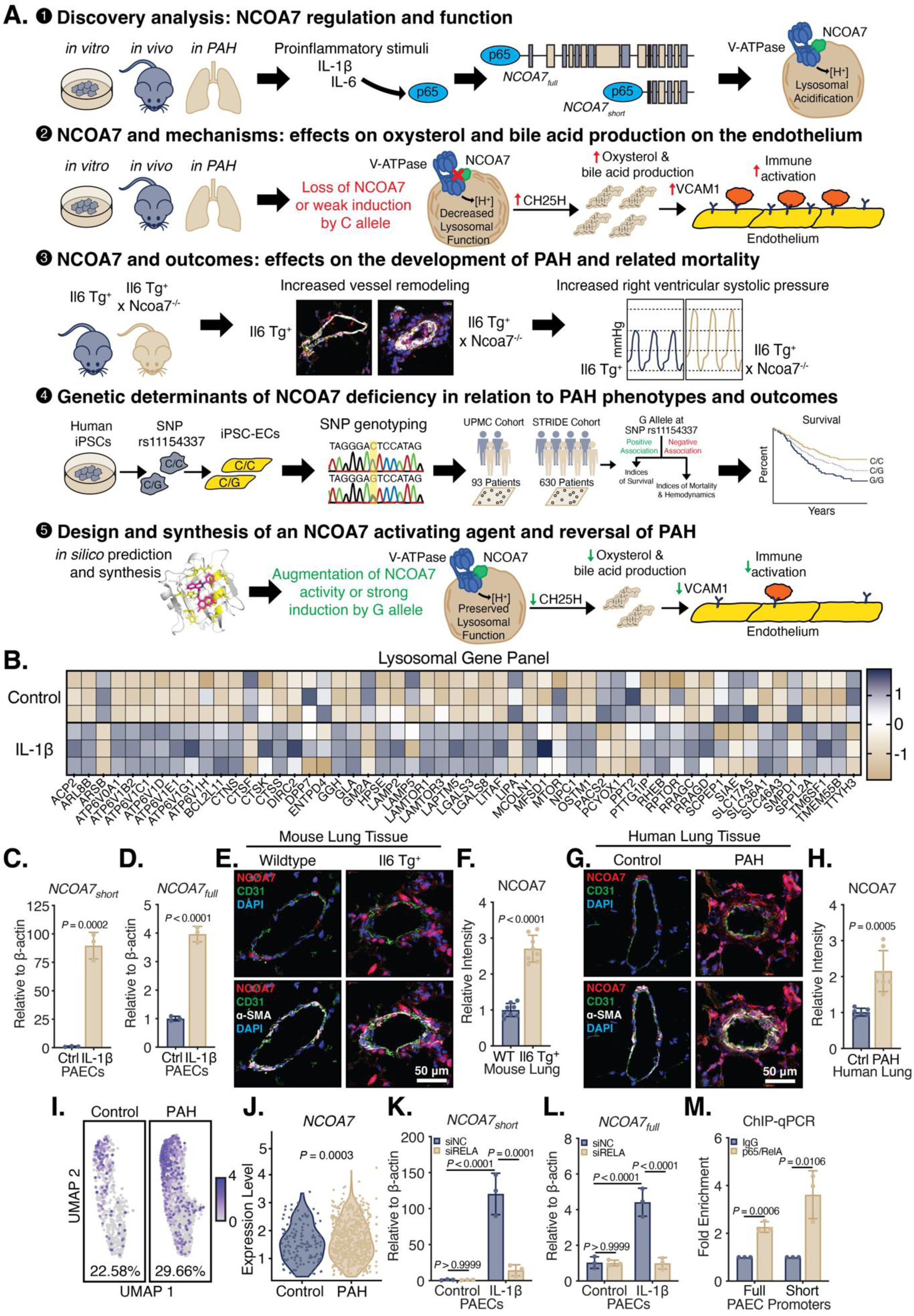
Convergent inflammatory regulation of NCOA7 across cellular, animal, and human instances of PAH. (**A**) Roadmap summary of the study. (**B**) Transcriptomic analysis of human PAECs under control or IL-1β (N=3/group). Z-score presented as positive in blue and negative in gold. Genes listed have an FDR-corrected *P*-value < 0.05. (**C** and **D**) *NCOA7* isoform expression via RT-qPCR (N=3/group). (**E** and **F**) Immunofluorescent (IF) staining for and (**G** and **H**) quantification of NCOA7 (red), CD31^+^ ECs (green), α-SMA^+^ smooth muscle cells (white), and DAPI-stained nuclei (blue) in pulmonary vessels of mouse (N=8/group) and human PH models (N=6-7/group). (**I** and **J**) violin plots of ECs expressing *NCOA7* identified via single-cell RNA sequencing from lungs of control or PAH patients (N>3/group). Cells were identified as expressing *NCOA7* if the transformed expression value was > 0.2. (**K** and **L**) *NCOA7* isoform expression under RNAi against *RELA* (N=3/group). Two-way ANOVA. (**M**) ChIP-qPCR against p65/RelA binding to full- and short-length promoter regions (N=3/group). All data are analyzed by Student’s *t*-test unless otherwise specified and presented as mean ± standard deviation.

### Discovery analyses: convergent inflammatory regulation of NCOA7 across cellular, animal, and human instances of PAH

Given the association between lysosomal dysfunction and oxysterol and bile acid production (*27*), we first sought to determine if triggers of EC dysfunction and PAH control lysosomal behavior. To do so, an unbiased, transcriptomic analysis was performed on primary human pulmonary artery endothelial cells (PAECs) exposed to IL-1β—a proinflammatory cytokine elevated in the plasma of PAH patients and known trigger of disease pathogenesis (*35*). Lysosomal regulatory genes were globally upregulated with a distinct subset comprising V-ATPase subunits, which are binding partners of NCOA7 that promote acidification of the lysosome (*17–19*) (**Fig. 1B**). Correspondingly, IL-1β upregulated NCOA7—both the canonical, full-length isoform (*NCOA7_full_*) and, to a greater extent, an alternative-start, short-length version of *NCOA7* (*NCOA7_short_*) (**Figs. 1C and 1D, fig. S1A**).

Other triggers of EC dysfunction in PAH similarly upregulated *NCOA7*. Specifically, exposure to the proinflammatory cytokine IL6 and its soluble receptor (IL6Rα), which has been linked to PAH (*35*), induced both short- and full-length isoforms (**figs. S1B and S1C**). Hypoxia, a well-established driver of PH, (*7*) also increased both isoforms (**figs. S1D and S1E**). Taken together, these data indicate a potential role for *NCOA7*, especially its unique, alternative-start isoform, across multiple triggers of PH.

In a severe inflammatory rodent model of PAH, transgenic mice under constitutive IL-6 overexpression and chronic hypoxia demonstrated elevated *Ncoa7* expression in CD31^+^ ECs isolated from lung tissue (**figs. S1F and S1G**). This was also observed in the pulmonary vessels of *Il6* transgenic mice without hypoxia—a milder form of experimental PAH (**Figs. 1E and 1F**). Similarly, examining the *in situ* localization of NCOA7, there was marked and transmural upregulation in pulmonary vessels with a heightened expression localized to the endothelium—in both the chronic hypoxia mouse model and the monocrotaline-exposed PAH rat model (**figs. S1H–S1K**).

In human lung tissue from patients with Group I PAH, NCOA7 expression was elevated within pulmonary vessels when compared to healthy controls (**Figs. 1G and 1H**). Single-cell RNA sequencing performed on lungs from idiopathic Group I PAH patients revealed an increased number of ECs expressing *NCOA7* when compared to healthy controls (22.58% versus 29.66%) (**Fig. 1I**) (*24*). Moreover, in NCOA7-positive ECs, *NCOA7* expression was upregulated in PAH patients (**Fig. 1J**).

Thus, in proinflammatory models of PH using primary PAECs, rodent models, and human patients, we found that NCOA7 was upregulated within the pulmonary vessel and, most importantly, the endothelium. However, since NCOA7 deficiency and consequent loss of lysosomal acidification should increase inflammation and worsen disease in other contexts (*10*), we hypothesized that NCOA7 acts as a homeostatic brake under proinflammatory stress to reduce disease pathogenesis through attenuation of EC immunoactivation.

To investigate upstream inflammatory mechanisms that modulate *NCOA7*, binding sites for the well-established inflammatory transcription factor complex—the RelA/p65 (*RELA*) subunit of NF-κB—were predicted in the canonical (*i.e.*, full-length) and non-canonical (*i.e.*, short-length) *NCOA7* promoter regions (*36*). Correspondingly, *RELA* knockdown in PAECs abrogated the IL-1β-mediated upregulation of both isoforms (**Figs. 1K and 1L**). A chromatin immunoprecipitation coupled with quantitative PCR (ChIP-qPCR) revealed significant enrichment of RelA/p65 at DNA sequences of the canonical and non-canonical promoters (**Fig. 1M**), indicating direct promoter-transcription factor binding.

### Loss of NCOA7 and mechanisms: lysosomal dysfunction and lipid accumulation

To investigate a putative link between NCOA7 and oxysterol production in the presence of proinflammatory conditions, we first characterized the NCOA7-mediated control of lysosomal acidification, given the known function of lysosomes in sterol trafficking (*37, 38*). In human PAECs exposed to IL-1β, knockdown of *NCOA7* reversed the interleukin-specific alteration of network of genes governing lysosomal function (**Fig. 2A**). Notably, a number of these genes (*e.g.*, *ATP6V0A1*, *ATP6V1B2*, *ATP6V1C1*, *ATP6V1D*, *ATP6V1E1*, *ATP6V1G1*, and *ATP6V1H*) encode for subunits of V-ATPases—machinery necessary for lysosomal acidification and thus the function of pH-sensitive enzymes.

**Fig. 2.**
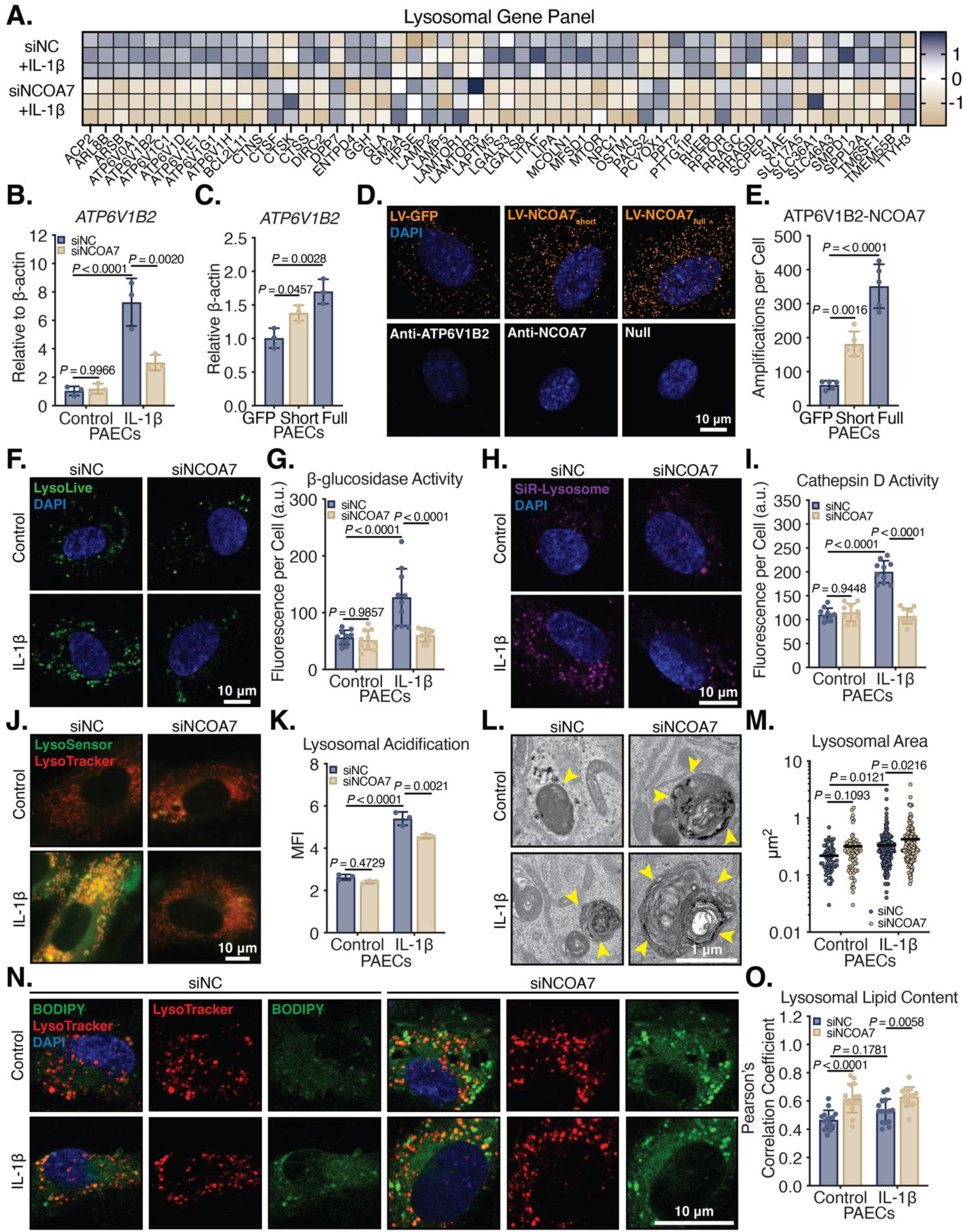
NCOA7 deficiency results in lysosomal dysfunction and lipid accumulation under proinflammatory conditions. (**A**) Transcriptomic analysis of PAECs under IL-1β subjected to RNAi against control or *NCOA7* (N=3/group). Z-score presented as positive in blue and negative in gold. Identified lysosomal genes have an FDR-corrected P-value < 0.05. (**B**) Expression of *ATP6V1B2* under siNC or siNCOA7 via RT-qPCR (N=3/group). (**C**) Expression of *ATP6V1B2* with lentiviral delivery of control (LV-GFP), NCOA7_short_, or NCOA7_full_ isoforms (N=3/group). Data analyzed by one-way ANOVA. (**D** and **E**) Association of the V-ATPase subunit ATP6V1B2 with NCOA7 measured by proximity ligation assay (orange). Bottom panel demonstrates control images of ATP6V1B2, NCOA7, or neither antibody. Top panel demonstrates dual incubation of ATP6V1B2 and NCOA7 antibodies with lentiviral transduction of GFP control, NCOA7_short_, or NCOA7_full_. Quantification of amplified signal per cell (N=5 cells/group). Data analyzed by one-way ANOVA. (**F** and **G**) LysoLive probe (green) reflecting β-galactosidase activity and thus lysosomal acidification. Quantified as fluorescence per cell (N=10 cells/group). (**H** and **I**) Silicone rhodamine (SiR)-Lysosome dye targeting active cathepsin D (purple) and indicating low lysosomal pH. Quantified as fluorescence per cell (N=10 cells/group). (**J**) Live-cell confocal microscopy of lysosomes (LysoTracker Deep Red) and lysosomal acidification (LysoSensor Green DND-189). (**K**) Median fluorescence intensity (MFI) ratio of LysoSensor Yellow/Blue DND-160 dye via flow cytometry representing lysosomal acidification (N=3/group). (**L** and **M**) Representative images and quantification of lysosomal area in transmission electron micrographs (N=20/group). Yellow arrows indicate lamellar-like inclusions reflecting lipid accumulation within lysosomes. (**N** and **O**) IF staining of colocalization (yellow) of neutral sterols (BOPIDY in green) and acidic lysosomes (LysoTracker in red). Colocalization was measured using EzColocalization and plotted as Pearson’s Correlation Coefficient (N=15 cells/group). All data are analyzed by two-way ANOVA unless otherwise specified and presented as mean ± standard deviation.

Previous proteomics-based studies established that NCOA7 interacts with ATP6V1B1— a renal specific paralog of ATP6V1B2 (*19, 39, 40*). Correspondingly, in PAECs, *NCOA7* knockdown abrogated the IL-1β-mediated upregulation of *ATP6V1B2*, and the forced overexpression of either the short- or full-length isoforms upregulated *ATP6V1B2* (**Figs. 2B and 2C, figs. S1L–S1P**). In addition, ATP6V1B2 was upregulated in the pulmonary endothelium of rodent and human models of PH (**fig. S2A–S2H**). To assess for a direct interaction between these proteins, a proximity ligation assay demonstrated perinuclear staining indicative of ATP6V1B2-NCOA7 interactions and consistent with a perinuclear distribution of lysosomes (*41*). Moreover, forced overexpression of NCOA7_short_ or NCOA7_full_ upregulated the number of ATP6V1B2-NCOA7 interactions in the lysosome (**Figs. 2D and 2E**). These data demonstrated a role for both short- and full-length NCOA7 as regulatory components of the V-ATPase complex with putative downstream lysosomal function.

To assess NCOA7 activity in modulating lysosomal acidification, two quantitative measures of lysosomal enzyme activity were utilized. First, cleavage of the fluorescent LysoLive tracer (*42*) was increased by IL-1β and subsequently blocked with loss of *NCOA7* (**Figs. 2F and 2G**). Second, lysosome-dependent cathepsin D activity was assessed utilizing SiR-Lysosome (*43*). In line with the IL-1β-mediated increase of *ATP6V1B2*, cathepsin D activity was significantly upregulated by IL-1β, as noted by SiR-Lysosome fluorescence, and was abrogated by *NCOA7* knockdown (**Figs. 2H and 2I**).

To directly evaluate lysosomal acidification, the acidotropic probe LysoSensor Green DND-189 was utilized for its accumulation in acidic compartments and enhanced fluorescence under acidic conditions. Consistent with the IL-1β-mediated increase in lysosomal enzyme activity, IL-1β increased the LysoSensor fluorescent signal, which was reversed with *NCOA7* knockdown (**Fig. 2J**). Additionally, IL-1β drove a shift to yellow fluorescence in PAECs when using the acidotropic probe LysoSensor Yellow/Blue DND-160 (*44*), indicating enhanced acidification of the lysosomal lumen (**Fig. 2K**). The addition of NCOA7 deficiency reversed the IL-1β-mediated yellow fluorescence shift. These observations mimic findings in LSDs, which are notable for the accumulation of undigested cellular components in the lysosomal compartment (*45*).

Failure of such V-ATPase complex formation or lysosomal acidification is a known driver of lysosomal dysmorphology (*46*). Accordingly, morphologic analysis of lysosomes using transmission electron microscopy of human PAECs deficient for NCOA7 revealed marked hypertrophy of lysosomes as quantified by lysosomal area (**Figs. 2L and 2M**), denoting an inability of the lysosomal compartment to breakdown and process cellular components. Moreover, the enlarged lysosomes in NCOA7-deficient cells carried lamellar-like inclusions (**Figs. 2L and 2M**; yellow arrows), indicative of sterol buildup—again phenocopying LSDs that present with abnormal lysosomal sterol accumulation (*27*).

To address whether the lamellar-like structures observed in the electron micrographs were lipids, PAECs were co-stained using a dye against neutral lipids (*i.e.*, BODIPY) and a dye that specifically localizes to acidic compartments (*i.e.*, LysoTracker). With loss of NCOA7, there was an increase in lipid punctae throughout the cell, which specified vesicular accumulation.

Correspondingly, the hyperintense lipid punctae were identified within acidic vesicles, supporting the idea that loss of NCOA7 caused such lysosomal accumulation of lipids (**Figs. 2N and 2O**). Taken together, these findings establish NCOA7 as a binding partner to the V-ATPase complex to promote lysosomal acidification and sterol trafficking in PAECs.

### Loss of NCOA7 and mechanisms: reprogramming of sterol metabolism via abnormal lipid accumulation

Alterations in lysosomal trafficking affect sterol homeostasis (*47*). Correspondingly, transcriptomic analysis of *NCOA7*-deficient human PAECs revealed marked enrichment and downregulation of biosynthetic processes related to sterol metabolism (**Figs. 3A and 3B**; red arrows). In a sterol saturated cell, uptake pathways of extracellular cholesterol are inhibited, specifically through reduction of the low-density lipoprotein receptor (LDLR) density on the cell membrane (*48*). *NCOA7* deficiency in PAECs reduced *LDLR* expression (**Fig. 3C**), accompanied by a functional attenuation in the uptake of fluorescently labeled cholesterol (**Figs. 3D and 3E**). Total cholesterol content in *NCOA7*-deficient PAECs was also upregulated, while forced overexpression of NCOA7 reduced total cholesterol content (**Figs. 3F and 3G**). No appreciable differences were detected in post-squalene intermediates in NCOA7-deficient versus NCOA7-replete PAECs (**figs. S3A–S3N**), indicating that NCOA7-dependent modulation of sterol intermediate flux does not rely upon *de novo* cholesterol synthesis. Thus, the downregulation of sterol metabolism by NCOA7 deficiency is driven primarily by lysosomal alterations of sterol handling rather than by *de novo* synthesis.

**Fig. 3.**
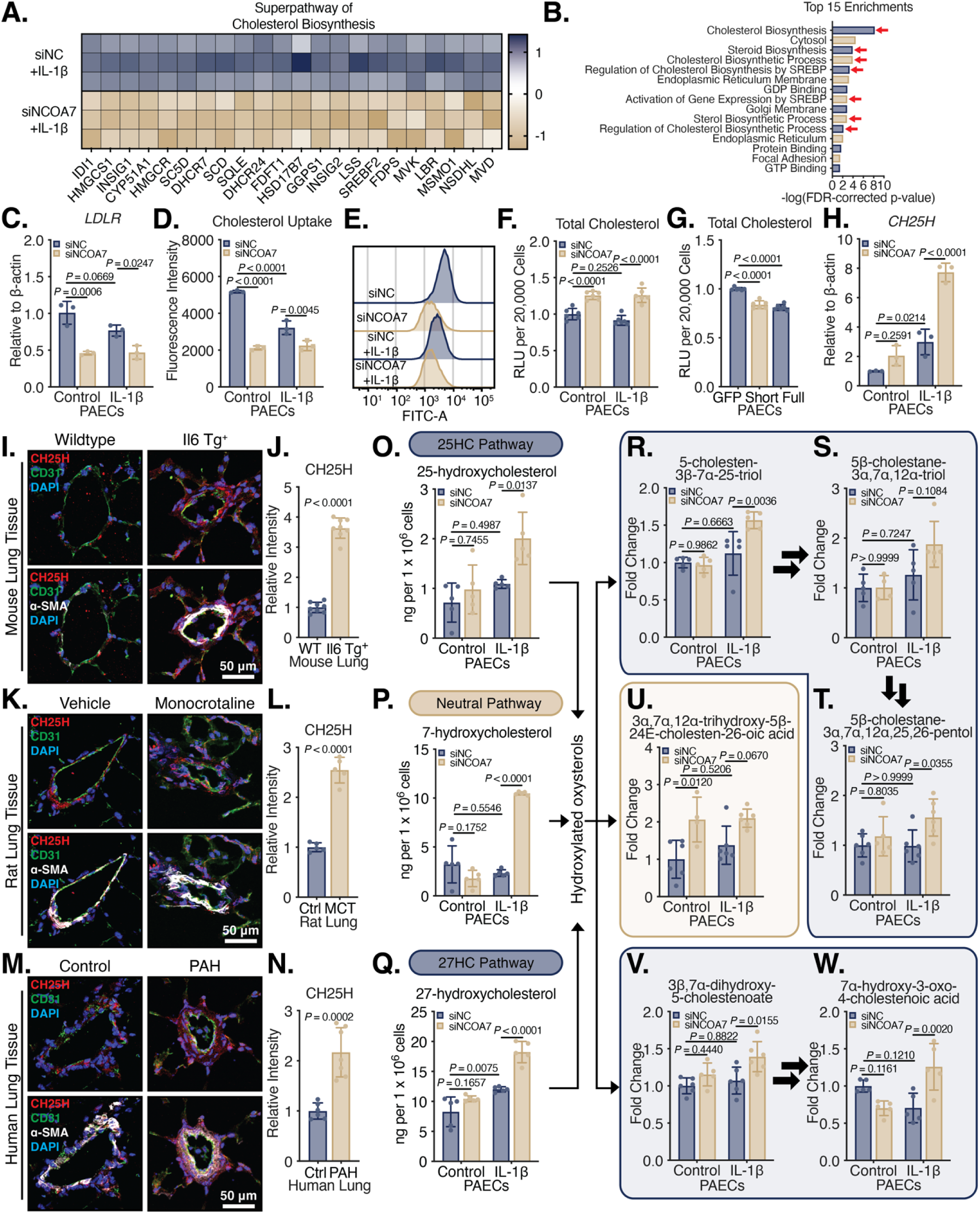
NCOA7 deficiency reprograms sterol metabolism to upregulate oxysterols and bile acids. (**A**) Transcriptomic analysis of PAECs under IL-1β subjected to RNAi against control or *NCOA7* (N=3/group). Z-score presented as positive in blue and negative in gold. Identified cholesterol metabolism genes have an FDR-corrected P-value < 0.05. (**B**) Gene set enrichment analysis of top 15 pathways by FDR-adjusted p-value with a majority related to sterol metabolism and homeostasis (highlighted with red arrows). (**C**) Expression of *LDLR* in PAECs under IL-1β subjected to RNAi against control or *NCOA7* (N=3/group). (**D** and **E**) Flow cytometric analysis of fluorescently tagged NBD-cholesterol uptake (N=3/group). (**F**) Total cholesterol content as measured by relative luminescence under siNC or siNCOA7 or (**G**) with overexpression using lentiviral GFP, NCOA7_short_, or NCOA7_full_. Data analyzed by one-way ANOVA (N=6/group). (**H**) Expression of *CH25H* in PAECs under IL-1β subjected to RNAi against control or *NCOA7* via RT-qPCR (N=3/group). (**I** to **N**) IF staining for and quantification of CH25H (red), CD31^+^ ECs (green), α-SMA^+^ smooth muscle cells (white), and DAPI-stained nuclei (blue) in the pulmonary vessels of mouse (N=8/group), rats (N=5/group), and human (N=6-7/group) PH models. Student’s *t*-test. (**O** to **Q**) Targeted oxysterol quantification via LC-MS (N=5/group) and (**R** to **W**) unbiased bile acid metabolite quantification via LC-MS organized by proposed pathways in shaded boxes (N=4-6/group). The metabolite 7HOCA is depicted in panel (**W**). All data are analyzed by two-way ANOVA unless otherwise specified and presented as mean ± standard deviation.

To protect against cholesterol accumulation, the cell can engage either in its direct export through transporters or by increasing its solubility through a series of oxidative steps. Accordingly, deficiency of *NCOA7* significantly upregulated cholesterol 25-hydroxylase (*CH25H*)—an oxysterol-generating enzyme that increases cholesterol solubility (**Fig. 3H**). Revealing the *in vivo* relevance of these processes, CH25H was upregulated in the pulmonary vessels of proinflammatory rodent models of PAH and Group I PAH patients with localization to the endothelium (**Figs. 3I−3N and figs. S3O and S3P**). Overall, these data established a central role for NCOA7 in the maintenance of EC sterol homeostasis through the generation of oxidized sterol species in cultured PAECs and in diseased endothelium *in vivo*.

### Loss of NCOA7 and mechanisms: induced endothelial generation of oxysterols and downstream bile acid derivatives

To corroborate whether the observed upregulation of *CH25H* enhanced oxysterol production, we performed targeted lipidomic analysis using liquid chromatography-mass spectrometry (LC-MS). *NCOA7* knockdown in human PAECs exposed to IL-1β significantly upregulated 25-hydroxycholesterol (25HC), 27-hydroxycholesterol (27HC), and autoxidation-generated 7α-hydroxycholesterol (**Figs. 3O−Q**). These oxysterols are known to be metabolized into downstream bile acid derivatives through incompletely understood mechanisms (*49*). Accordingly, *NCOA7* knockdown upregulated several downstream bile acid derivatives (*34*) in sequential pathways, such as 5-cholesten-3β-7α,25-triol, 5β-cholestane-3α,7α,12α-triol, and 5β-cholestane-3α,7α,12α,25,26-pentol (**Figs. 3R−3T**) and the 7α-hydroxycholesterol derivative 3α,7α,12α-trihydroxy-5β-24E-cholesten-26-oic acid (**Fig. 3U**). Moreover, the upstream metabolites 3β,7α-dihydroxy-5-cholestenoate and 7α-hydroxy-3-oxo-4-cholestenoic acid (7HOCA) were also upregulated (**Figs. 3V and 3W**). Thus, consistent with the upregulation of the oxysterol-generating enzyme CH25H (**Fig. 3H**), NCOA7 deficiency induced production of the numerous oxidized cholesterol metabolites and downstream bile acids in ECs.

### Loss of NCOA7 and mechanisms: endothelial dysfunction through oxysterol generation

Given the immunomodulatory functions of oxysterols in diseased endothelium (*50*), we sought to determine if NCOA7 deficiency relied upon oxysterols to promote EC dysfunction. NCOA7 deficiency further upregulated the vascular cellular adhesion molecule 1 (*VCAM1*)—a surrogate of endothelium immunoactivation (**Figs. 4A and 4B and fig. S4P**)—when under proinflammatory stimulation with IL-1β. Conversely, forced overexpression of NCOA7 isoforms reversed VCAM1 expression (**Figs. 4C and 4D and fig. S4Q**). To determine if *NCOA7* deficiency was dependent upon downstream, oxidized forms of cholesterol to induce PAEC pathophenotypes, concomitant knockdown experiments against the oxysterol generating enzyme CH25H, which is upregulated with *NCOA7* deficiency, were performed. Notably, inhibition of CH25H induction under NCOA7 deficiency prevented VCAM1 expression (**Figs. 4E and 4F and fig. S4R**).

**Fig. 4.**
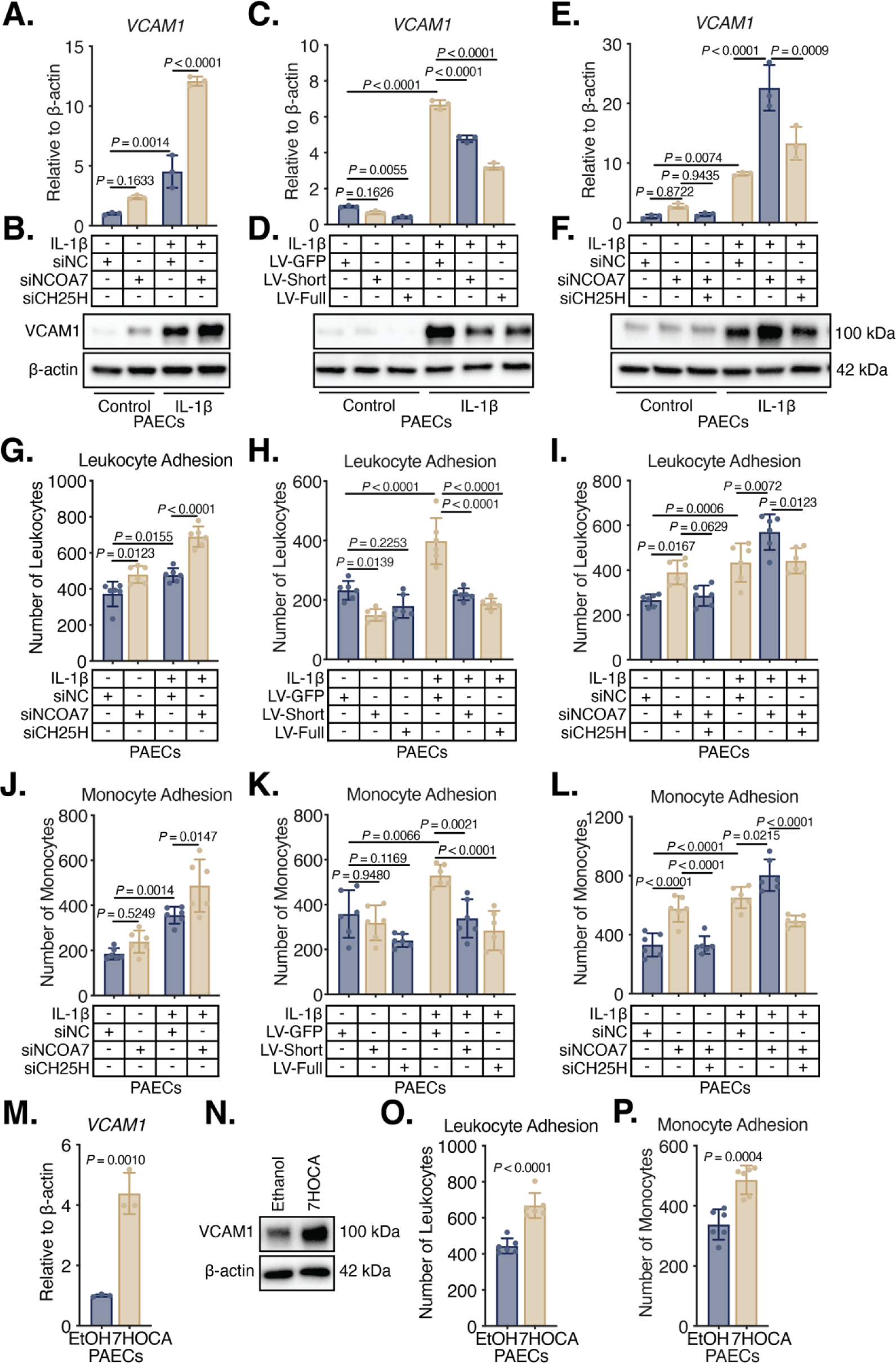
The NCOA7-CH25H axis drives pulmonary endothelial immunoactivation. (**A** to **F**) VCAM1 expression via RT-qPCR and immunoblot under (**A** and **B**) RNAi against *NCOA7*, (**C** and **D**) lentiviral delivery of *NCOA7_short_* or *NCOA7_full_*, or (**E** and **F**) RNAi against *NCOA7* and *CH25H* (N=3/group). Immune cell adhesion of leukocytes (**G** to **I**) or monocytes (**J** to **L**) to an endothelial monolayer (N=6/group). (**M** and **N**) Expression of VCAM1 by RT-qPCR and immunoblot in PAECs treated with ethanol control versus 7HOCA (50 μM) for 24 hours (N=3/group; Student’s *t*-test). (**O** and **P**) Leukocyte and monocyte adhesion in 7HOCA-treated PAECs compared to ethanol controls (N=6/group; Student’s *t*-test). All data are analyzed by two-way ANOVA unless otherwise specified and presented as mean ± standard deviation.

In line with observed VCAM1 changes under various states of NCOA7 and CH25H expression, we observed increased attachment of both leukocytes and monocytes to an EC monolayer deficient for NCOA7 (**Figs. 4G and 4J**), which could be reversed with NCOA7 overexpression (**Figs. 4H and 4K**), and a significant attenuation of immune cell attachment with inhibition of CH25H under conditions of NCOA7 deficiency (**Figs. 4I and 4L**).

In addition and consistent with the concept of an immunoactivated, apoptosis-resistant, and hyperproliferative endothelium in pulmonary vascular disease (*51, 52*), NCOA7 deficiency in PAECs abrogated IL-1β-mediated apoptosis while simultaneously enhancing proliferative capacity (**figs. S4A and S4D**). NCOA7 facilitated PAEC apoptosis under proinflammatory conditions (**fig. S4B**) and, in parallel, attenuated proliferation with more pronounced inhibition under IL-1β (**fig. S4E**). Inhibition of *CH25H* upregulation, however, reversed the attenuation of apoptosis, while enhancing the proliferative capacity of the cells (**figs. S4C and S4F**). In sum, the presence of NCOA7 acted as a homeostatic brake under proinflammatory stimuli, preventing immunoactivation of the endothelium with induction of apoptosis and inhibition of proliferation.

To determine if bile acids are sufficient to immunoactivate the endothelium, 7HOCA was directly applied to PAECs in culture. Displaying its proinflammatory nature, 7HOCA significantly upregulated VCAM1 (**Figs. 4M and 4N and fig. S4S**), thus promoting adhesion of both leukocytes and monocytes to a PAEC monolayer (**Figs. 4O and 4P**). Notably, the direct application of 25HC and downstream derivatives, such as a triol and tetrol, similarly upregulated VCAM1 and enhanced immune cell adhesion to a PAEC monolayer (**figs. S4G−S4O**). In sum, these findings demonstrate that oxysterol-generating enzymes and their downstream oxidized sterol species are necessary and sufficient in mediating immune activation of the NCOA7-deficient pulmonary endothelium.

### Loss of NCOA7 and outcomes: orotracheal delivery of 7HOCA worsens PAH in vivo

To determine if the presence of NCOA7 protects against EC immunoactivation in PAH severity, we utilized an *Il6* transgenic (Tg^+^) mouse to elicit severe pulmonary inflammation as a model of angioproliferative PAH (*53*). A whole-body knockout mouse for *Ncoa7* was crossed onto *Il6* Tg^+^ mice to determine if loss of NCOA7 would worsen indices of PH *in vivo* (**Fig. 5A**). Echocardiographic assessment excluded any gross alterations in heart rate and left ventricular function, as noted by left ventricular fractional shortening (LVFS), left ventricular ejection fraction (LVEF), and left ventricular posterior wall distance during diastole and systole (LVPW;d and LVPW;s) (**figs. S5A−S5E**).

**Fig. 5.**
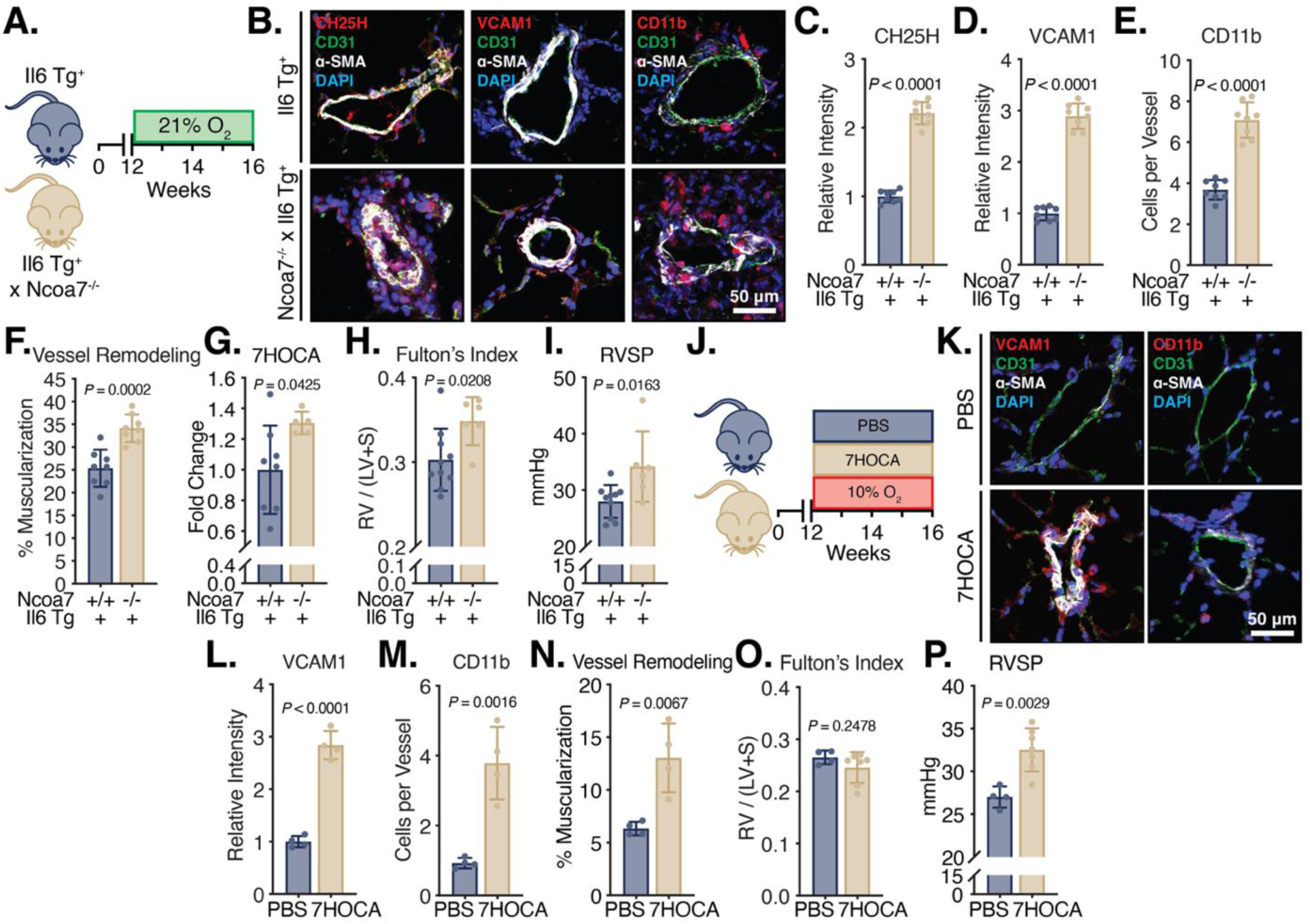
Genetic loss of *NCOA7* and the orotracheal delivery of 7HOCA worsens PAH *in vivo*. (**A**) *Ncoa7*-null mice crossed onto the *Il6* Tg^+^ PAH model. (**B** to **F**) Pulmonary vessels from *Il6* Tg^+^ versus *Ncoa7^-/-^*x *Il6* Tg^+^ mice stained for a target protein (*i.e.*, CH25H, VCAM1, or CD11b; red), the endothelial layer (CD31; green), the smooth muscle layer (α-SMA; white), and nuclear counterstain (DAPI; blue). Quantification of the relative intensity of CH25H or VCAM1, the number of CD11b^+^ cells per vessel, or the degree of vessel muscularization defined by α-SMA layer thickness to total vessel diameter. N=8/group. (**G**) Measurement of 7HOCA using LC-MS in the serum of *Il6* Tg^+^ versus *Ncoa7^-/-^* x *Il6* Tg^+^ mice (N=5-8/group). (**H** and **I**) Fulton’s Index and RVSP of *Il6* Tg^+^ versus *Ncoa7^-/-^* x *Il6* Tg^+^ mice (N=6-10/group). (**J**) Mice received orotracheal delivery of either PBS or 7HOCA (10 mg/kg) for four weeks under hypoxic (10% O_2_) conditions and sacrificed at 16 weeks. (**K** to **N**) Pulmonary vessels from PBS versus 7HOCA mice stained for a target protein (*i.e.*, VCAM1, or CD11b; red), the endothelial layer (CD31; green), the smooth muscle layer (α-SMA; white), and nuclear counterstain (DAPI; blue). Quantification of the relative intensity of VCAM, the number of CD11b^+^ cells per vessel, or the degree of vessel muscularization defined by α-SMA layer thickness to total vessel diameter (N=4/group). (**O** and **P**) Fulton’s Index and RVSP of mice receiving orotracheal PBS or 7HOCA (N=4-7/group). All data are analyzed by Student’s *t*-test unless otherwise specified and presented as mean ± standard deviation.

*Ncoa7*-null mice displayed elevated CH25H expression in pulmonary arterioles, accompanied by elevated plasma levels of 7HOCA and downstream tetrol species of 25HC (**Figs. 5B, 5C, 5G, and figs. S5F–S5I**). These findings corresponded with the oxysterol and bile acid plasma signatures associated with PAH severity in humans (*34*) and our studies of cultured PAECs. The elevation of 7HOCA in *Ncoa7*-deficient mice resulted in immunoactivation of the endothelium as noted by enhanced VCAM1 expression and CD11b^+^ monocyte infiltration **(Figs. 5B, 5D, and 5E**). Moreover, *Ncoa7*-null mice displayed increased pulmonary arteriole muscularization (**Fig. 5F**), accompanied by worsened hemodynamic manifestations of PAH with increased right ventricular systolic pressure (RVSP) and Fulton index—a measure of right ventricular remodeling (**Figs. 5H and 5I**).

Finally, to assess the pathogenicity of 7HOCA directly, we utilized a chronically hypoxic mouse model with serial, orotracheal deliveries of either normal saline or 7HOCA (**Fig. 5J**) and confirmed via echocardiography no alterations in left ventricular function **(figs. S5J–S5N**). As expected, the orotracheal delivery of 7HOCA upregulated immunoactivation of the endothelium, as noted by enhanced VCAM1 expression and CD11b^+^ monocyte infiltration into the vascular bed **(Figs. 5K−5M**). Consistent with the genetic knockout of *Ncoa7*, 7HOCA worsened PAH, as reflected by increased pulmonary arteriolar remodeling and increased RVSP despite an unchanging Fulton’s Index (**Figs. 5N−5P**). Taken together, our data demonstrated that either a genetic loss of *Ncoa7* or direct delivery of the proinflammatory sterol 7HOCA were sufficient to promote PH *in vivo*.

### Loss of NCOA7 and outcomes: oxysterols and bile acids predict morbidity and mortality in clinical PAH

Underscoring the clinical importance of this mechanism in controlling disease severity, we and our colleagues identified a plasma signature inclusive of the same NCOA7-dependent sterols and bile acids associated with PAH mortality (adjusted *P*<1.1x10^-6^). In our parallel study (*34*), via an unbiased, metabolome-wide association study from the multicenter PAH Biobank cohort (N=2,756), Alotaibi et *al*. identified 13 distinct plasma oxysterols and bile acids that best predicted four-year mortality in PAH (AUC 73%, 95% CI 68% to 78%). Notably, of these top 13 oxysterols and bile acids, four were the same metabolites upregulated in ECs deficient for *NCOA7* (**Figs. 3Q, 3R, 3S, and 3W**) and in the serum of *Ncoa7*-deficient mice (**Fig. 5G** and **figs. S5F−S5I**). These findings emphasize the clinical importance of NCOA7-dependent oxysterols and bile acids in controlling PAH severity, thereby establishing a framework that links lysosome-dependent inflammation with both population-level and metabolome-wide signals in PAH.

### Intronic SNP rs11154337 controls RelA/p65 binding to the non-canonical promoter of NCOA7 and is associated with PAH disease severity and mortality

Our findings thus far define NCOA7 as a homeostatic brake in proinflammatory conditions that preserves lysosomal activity and sterol trafficking to attenuate EC immunoactivation in PAH. Based on the role of single nucleotide polymorphisms (SNPs) in regulating gene promoter activity (*54*), we sought to determine if pathogenic NCOA7 deficiencies could result from genetic, SNP-dependent control of NCOA7 expression and its downstream function (**Figs. 1–5**). First, we surveyed annotated SNPs based on their proximity to the canonical and non-canonical promoters and high levels of epigenetic marks indicating elevated transcriptional activity. In doing so, we found a candidate SNP—rs11154337 (GRCh 38, chr6:125922445)—located near an intronic region proximal to the non-canonical promoter of *NCOA7* and carrying a substantial burden of histone modifications (*36*). Second, given the tandem regulation of both short and long isoforms of NCOA7 in PAH (**Fig. 1**), we sought to discern any potential regulatory function of this SNP via positional backfolding onto the canonical promoter. To do so, we utilized publicly available high-throughput chromatin conformation capture (3C) on human umbilical vein endothelial cells (GEO IDs GSM3438650 and GSM3438651) from the 3D-genome Interaction Viewer & database (3DIV) (*55*). Such maps revealed that SNP rs11154337 carries long-range interactions greater than 120 kilobases with the canonical promoter region of *NCOA7* (**fig. S6A**). Chromosome conformation capture (3C) analysis performed on human PAECs demonstrated an interaction between the genomic loci encompassing SNP rs11154337 and the NCOA7 transcription start site (**figs. S6B and S6C**). Furthermore, ChIP against RelA/p65 revealed a significant enrichment for the SNP-containing region, supporting the presence of a p65 protein-SNP complex (**fig. S6D**).

Given that the SNP rs11154337 controls NCOA7 gene architecture and that NCOA7 controls the oxysterol signature associated with PAH severity and mortality (*34*), we sought to determine if this SNP was also associated with disease severity and mortality using two independent PAH cohorts. (**fig. S6E**). First, we analyzed a single-center PAH cohort of European-descent subjects from the University of Pittsburgh Medical Center with adjustments for age, sex, and vasodilator use (UPMC, **Table 1**, N=93). We found that the G allele was associated with a significant improvement of six-minute walk distance (*P*=0.0130; β=66.90, 95% CI [14.45-119.36]), **Fig. 6A**). Importantly, survival was significantly increased in patients who carried the homozygous G alleles (*P*=0.0250; hazard ratio=0.44, 95% CI [0.21-0.90], **Fig. 6B**). In a second, multicenter PAH cohort of European-descent from the Sitaxsentan To Relieve Impaired Exercise (STRIDE) trial comprising 45 United States and Canadian pulmonary hypertension centers (STRIDE, **Table 1**, N=630) (*56*), we validated that the presence of the G allele conferred a substantial survival benefit after adjustments for age, sex, PAH type, type of study inclusion, WHO functional classification, and vasodilator use (*P*=0.0002, hazard ratio=0.49, 95% CI [0.34–0.71], **Fig. 6C**). Thus, analysis of genomic, metabolomic, and clinical datasets across cohorts of PAH patients demonstrated the presence of interconnected activities between NCOA7 and SNP rs11154337 with glucuronidated oxysterols and clinical outcomes of PAH.

**Fig. 6.**
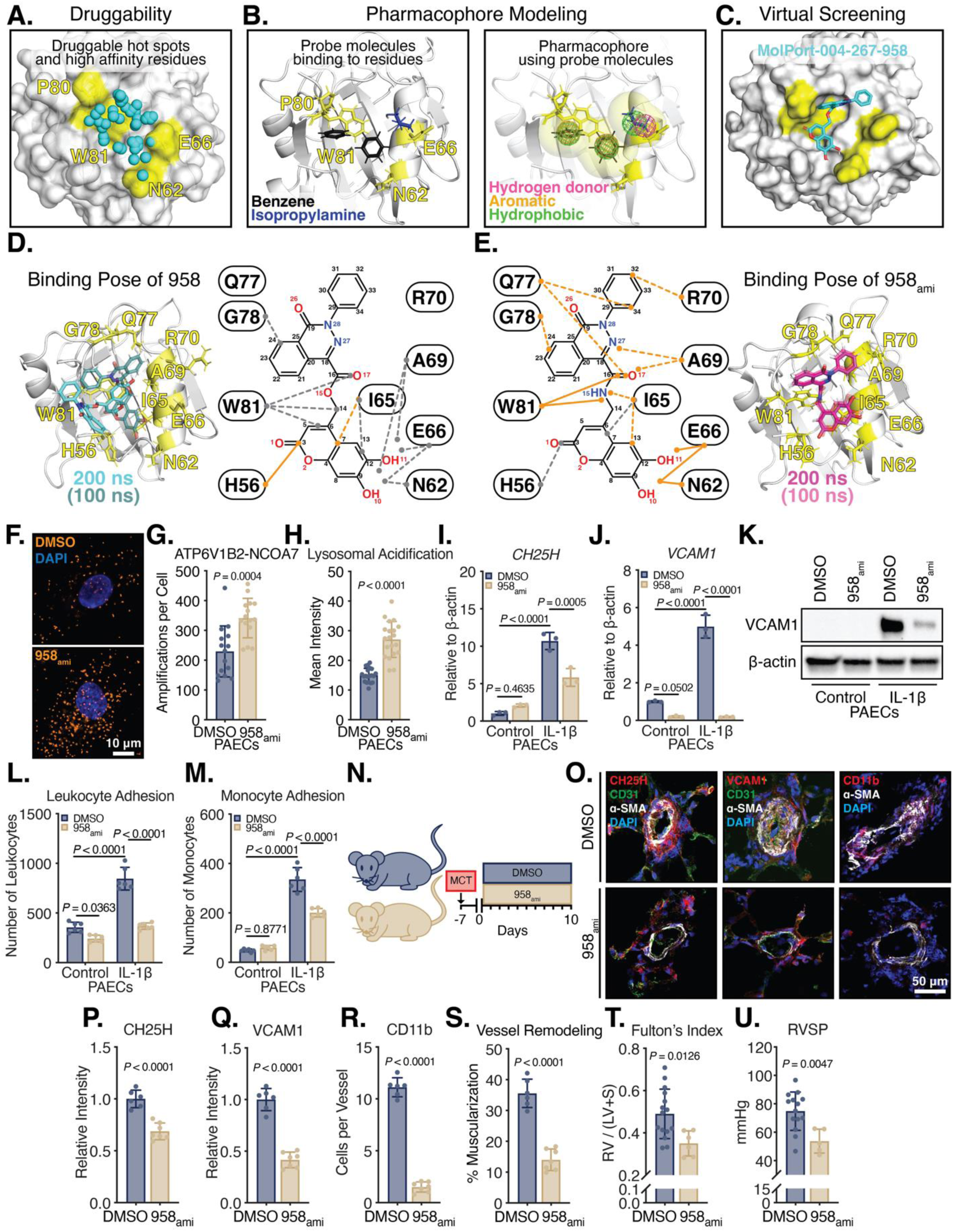
The G allele of SNP rs11154337 prevents lysosomal lipid accumulation and attenuates oxysterol-mediated immunoactivation in iPSC-ECs. (**A** to **C**) Allelic variants of SNP rs11154337 and their clinical readouts of six-minute walk distance (6MWD) and survival in PAH patients in the UPMC cohort (N=93) and STRIDE cohort (N=630). (**D**) Schematic of iPSC-EC production. (**E** and **F**) *NCOA7* isoform expression via RT-qPCR (N=3/group). (**G** and **H**) SiR-Lysosome dye against active cathepsin D (purple; N=10 cells/group). (**I**) *ATP6V1B2* expression via RT-qPCR (N=3/group). (**J**) Transmission electron micrograph quantification of lysosomal area. (**K**) Total cholesterol content as measured by relative luminescence (N=6/group). (**L** and **M**) BODIPY dye against neutral lipids (green; N=10 cells/group). (**N**) *CH25H* expression via RT-qPCR (N=3/group). (**O**) Targeted 25HC quantification via LC-MS (N=5/group). (**P** and **Q**) *VCAM1* expression via RT-qPCR or immunoblot (N=3/group). (**R** and **S**) Leukocyte and monocyte adhesion to iPSC-EC monolayer (N=6/group). All data are analyzed by two-way ANOVA unless otherwise specified and presented as mean ± standard deviation.

**Table 1.** Cohort demographics and clinical characteristics.

| **Demographic and clinical characteristic** | **Median [25% – 75%]** |
| --- | --- |
| **UPMC Cohort (N=93)** |  |
| Age at catheterization (years) | 51 [36 – 60] |
| Mean right atrial pressure (mmHg) | 9.00 [4.50 – 13.00] |
| Mean pulmonary artery pressure (mmHg) | 51.50 [40.75 – 60.25] |
| Pulmonary capillary wedge pressure (mmHg) | 10.00 [7.00 – 14.00] |
| Cardiac output (L/min) | 4.30 [3.45 – 5.40] |
| Cardiac index (L/min/kg/m<sup>2</sup>) | 2.30 [1.93 – 2.84] |
| Pulmonary vascular resistance (Wood units) | 9.70 [5.74 – 13.11] |
| **STRIDE Cohort (N=630)** |  |
| Age at catheterization (years) | 52.02 [38.98 – 62.68] |
| Mean right atrial pressure (mmHg) | 7.00 [5.00 – 11.00] |
| Mean pulmonary artery pressure (mmHg) | 50.00 [38.00 – 61.00] |
| Pulmonary capillary wedge pressure (mmHg) | 10.00 [7.00 – 13.00] |
| Cardiac output (L/min) | 4.30 [3.40 – 5.50] |
| Cardiac index (L/min/kg/m<sup>2</sup>) | 2.40 [1.96 – 3.00] |
| Pulmonary vascular resistance (Wood units) | 9.99 [6.00 – 14.00] |

### Genetic determinants of NCOA7: identification of an allele-specific SNP and its downstream pathogenic functions

Using this concept—and guided by the negative association of the G allele of the *NCOA7* intronic SNP rs11154337 to both the oxysterol signature predictive of mortality and clinical indices of PAH—we sought to determine if this SNP controls NCOA7 expression, lysosomal activity, and the production of oxysterol and bile acid metabolites to modulate EC behavior. To study the cellular and biological activity of SNP rs11154337 embedded near the non-canonical *NCOA7* promoter, we generated a set of genetically matched, isogenic inducible pluripotent stem cell (iPSC) lines with the allelic variants of SNP rs11154337 via CRISPR-Cas9-gene editing (C/C versus C/G genotypes, **Fig. 6D and fig. S6F**). The iPSCs were then differentiated into ECs (iPSC-ECs) and purified through a vascular endothelial cadherin (VE-Cadherin; also known as CD144)-based magnetic separation (*57*) (**fig. S6G**). Purified iPSC-ECs exhibited marked enrichment of the EC markers CD34, CD144, and CD309, and immunofluorescent staining of iPSC-ECs against CD144 and CD31 revealed a patterning consistent with endothelium (**figs. S6G and S6H**). Moreover, iPSC-ECs displayed angiogenic potential, as noted by vessel formation in growth factor-depleted Matrigel (**fig. S6H**).

C/G iPSC-ECs displayed higher expression of both short and long *NCOA7* isoforms when compared to the C/C line, confirming that the G allele increases *NCOA7* transcription (**Figs. 6E and 6F**). Providing a putative explanation for the long-range regulation of SNP rs11154337 and the canonical promoter of *NCOA7*, prior 3C data demonstrated a SNP rs11154337 interaction with the canonical promoter in human umbilical vein endothelial cells (**figs. S6A–S6D**) (*55*). Consistent with the observed differential in *NCOA7* expression and our prior findings with *NCOA7* knockdown, lysosomal activity, sterol homeostasis, and immunoactivation in iPSC-ECs were allele-dependent. iPSC-ECs carrying the G allele—and thus higher *NCOA7* expression— displayed a concomitant increase in its binding partner *ATP6V1B2* and subsequently lower lysosomal pH, as demonstrated by attenuated cleavage of SiR-Lysosome (**Figs. 6G−6I**). Moreover, the presence of the G allele prevented lysosomal hypertrophy in comparison to the iPSC-EC line homozygous for the C allele, indicating maintenance of proper lysosomal acidification and resultant sterol homeostasis (**Fig. 6J and fig. S6I**).

Similar to RNAi against *NCOA7*, less *NCOA7* expression in the homozygous C allele line resulted in higher sterol content and thus drove higher expression of *CH25H* and its metabolite 25HC (**Figs. 6K−6O**) that has implications for the generation of downstream oxidized species like 7HOCA. With greater production of 7HOCA, the homozygous C allele iPSC-line displayed elevated immunoactivation of the endothelium, reflected by enhanced VCAM1 expression and immune-cell adhesion (**Figs. 6P−6S and fig. S6J**). Thus, consistent with the association of the G allele of SNP rs11154337 as protective against oxysterol production and PAH severity, the G allele increased *NCOA7* expression, its downstream modulation of lysosomal acidification, oxysterol generation, and consequent EC immunoactivation.

### NCOA7 as therapeutic target: structural modeling and molecular simulations identify a novel activator of NCOA7

Toward identifying a small molecule activator of NCOA7, we performed structure-based computations composed of three parts: druggability simulations, pharmacophore modeling, and virtual screening (**Figs. 7A−7C**). Druggability simulations were carried out using the model structure of NCOA7 in the presence of explicit water and probe molecules representative of drug-like fragments (*58*). We used the probe molecules acetamide, acetate, benzene, imidazole, isobutane, isopropanol, and isopropylamine in six independent runs of 40 ns each. A molecular pocket was distinguished in three of the runs through its high affinity to bind the probe molecules (**Fig. 7A**, *cyan spheres*). This site also demonstrated hinge residues at or near the binding pocket from the Gaussian Network Model (GNM) analysis of NCOA7 (**fig. S7A**), including L83 (mode 1), L72 (mode 2), and E66 and W81 (mode 3) (**fig. S7B**). Notably, hinge sites have been shown in previous work to have a critical role in mediating the functional dynamics of proteins, and, as such, are used as target sites for binding small molecule modulators of protein function. For this reason, the identified molecular pocket was selected for further analysis using pharmacophore modeling boosted by both druggability simulations and GNM analysis.

**Fig. 7.**
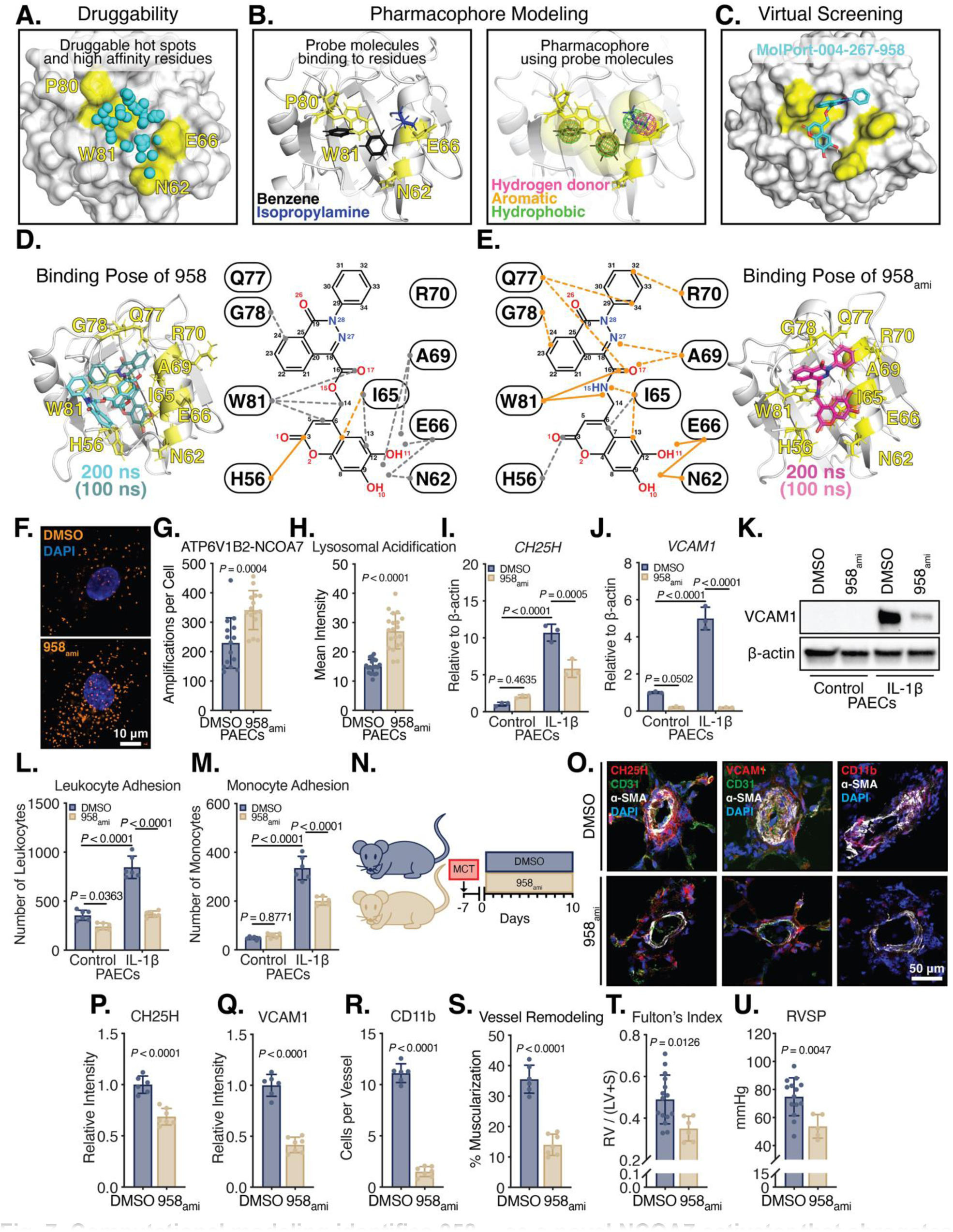
Computational modeling identifies 958_ami_ as a novel NCOA7 activator that abrogates endothelial immunoactivation and PAH. (**A** to **C**) Computational protocol for identifying small molecule modulators of NCOA7, comprised of three major components: (**A**) druggability simulations, (**B**) pharmacophore modeling, and (**C**) virtual screening. (**D** and **E**) Refinement of the identified compound 958 after MD simulations into its analogue 958_ami_. Compound atoms and NCOA7 residues interactions are shown in two dimensions. Stronger interactions are shown in orange dashed lines, while weaker interactions are shown in gray dashed lines. Orange solid lines are strong/persistent interactions with more than 0.3 µs cumulative duration per interaction over 0.6 µs total simulations. (**F** and **G**) Association of the V-ATPase subunit ATP6V1B2 with NCOA7 measured by proximity ligation assay (orange). Panels demonstrate dual incubation of ATP6V1B2 and NCOA7 antibodies with DMSO or 958_ami_. Quantification of amplified signal per cell (N=15 cells/group). (**H**) Lysosomal pH assessed via LysoSensor Green DND-189 and immunofluorescence intensity quantified per cell (N=20 cells/group). (**I**) *CH25H* expression via RT-qPCR (N=3/group; Two-way ANOVA). (**J** and **K**) Expression of VCAM1 by RT-qPCR and immunoblot in PAECs treated with DMSO versus 958_ami_ (20 μM) for 24 hours (N=3/group; Two-way ANOVA). (**L** and **M**) Leukocyte and monocyte adhesion in 958_ami_-treated PAECs compared to DMSO control (N=6/group. (**N**) Rats were loaded with monocrotaline (80 mg/kg i.p.) one week before initiation of a daily dose of 958_ami_ (7.5 mg/kg i.p.) for 10 days. (**O** to **S**) Pulmonary vessels from DMSO versus 958_ami_ injected rats stained for a target protein (*i.e.*, CH25H, VCAM1, or CD11b; red), the endothelial layer (CD31; green), the smooth muscle layer (α-SMA; white), and nuclear counterstain (DAPI; blue). Quantification of the relative intensity of CH25H or VCAM1, the number of CD11b^+^ cells per vessel, or the degree of vessel muscularization defined by α-SMA layer thickness to total vessel diameter. N=6-7/group. (**T**) Fulton’s Index and (**U**) RVSP of monocrotaline rats receiving intraperitoneal DMSO or 958_ami_. N=5-15/group. All data are analyzed by Student’s *t*-test unless otherwise specified and presented as mean ± standard deviation.

Using the tool Pharmmaker (*59*), we performed pharmacophore modeling, and subsequently selected the high affinity residues (*i.e.*, N62, E66, P80, and W81; **Fig. 7B**) at the identified molecular pocket. The interactions of these residues with the probe molecules were ranked based on their occurrence frequency during simulations. Notably, among the probe molecules, two benzenes demonstrated a high propensity to interact with N62, P80, and W81, while one isopropylamine interacted with E66 (**Fig. 7B**, *benzenes in black* and *isopropylamine in blue*). Molecular dynamics (MD) snapshots that simultaneously displayed multiple frequently observed (*i.e.,* entropically favored) interactions were used to construct the pharmacophore model, which was composed of one hydrogen bond donor, three hydrophobic rings, and two aromatic rings (**Fig. 7B**).

The pharmacophore model was then screened against the ZINC (*60*) and MolPort small molecule libraries via the Pharmit (*61*) server to obtain a ranked ensemble of compounds. The top scoring compounds were selected as hits for further experimental validation. Pilot screening suggested that compound (6,7-dihydroxy-2-oxo-2H-chromen-4-yl)methyl 4-oxo-3-phenyl-3,4-dihydrophthalazine-1-carboxylate (MolPort-004-267-958; herein called 958 in **Fig. 7C**) was an activator rather than an inhibitor of NCOA7. We then sought to further examine the binding behavior of 958 through molecular dynamics simulations.

### NCOA7 as therapeutic target: refinement of the activator 958 to optimize binding affinity

To create a molecule that more avidly binds to the molecular pocket of NCOA7, we performed MD simulations. As such, all-atom 0.6 µs MD simulations (or three independent, 0.2 µs runs) of 958 were performed to characterize the most critical functional groups and interactions (**Fig. 7C**). Based on these simulations, we designed an analogue of 958, where the O_15_ atom of the ester functional group is replaced with N_15_-H to create an amide functional group with the resultant compound herein called 958_ami_ (**Figs. 7D and 7E**). Subsequent MD simulations using 958_ami_ further clarified its enhanced activity. To compare the parental and analogue structures, we selected nine residues with high interaction affinities to the compounds: H56, N62, I65, E66, A69, R70, Q77, G78, and W81. Overall contact duration for these residues was greater than 0.35 µs out of 0.6 µs with 958 or 958_ami_. (**figs. S8A−S8C**). Strong interactions, defined as a contact duration greater than 0.3 µs, were formed by H56-O_1_ in 958 versus N62-O_10_, E66-O_10_/O_11_, and W81-N_15_/O_17_ in 958_ami_ (**Figs. 7D and 7E**; *orange solid lines*). Notably, atoms 16 through 34 do not undergo any significant interactions with the NCOA7 binding pocket; however, the substitution of O_15_ with N_15_-H resulted in multiple strong interactions with many residues within the pocket. Additionally, the substitution with N_15_ exhibited strong interactions with I65 and W81, caused new interactions of atoms 16 through 34 with A69, R70, and Q77, and strengthened the C_23_/C_24_-G78 and O_10_/O_11_-N62/E66 interactions. Lastly, using hydrogen bond analysis with a cutoff distance of 3.0 Å between donor and acceptor atoms with a 160° angle, E66 was identified with a significantly higher propensity to form hyrogen bonds with 958_ami_. Using PRODIGY-LIG (*62*), calculated binding affinities revealed that 958_ami_ was more stably bound to NCOA7 than the parental comound 958 (**fig. S9A**). The corresponding binding pose at 100 to 200 ns had a binding affinity of -8.16 ± 0.16 kcal/mol (**Fig. 7E**, *run 1*). 958_ami_ also had two additional stable poses: one where W81 lost its interaction with N_15_ (**fig. S9B**, *run 2*, -7.71 ± 0.16 kcal/mol) and the other where the compound rotated upside down within the pocket (**fig. S9B**, *run 3*, -8.83 ± 0.19 kcal/mol).

### NCOA7 as therapeutic target: administration of activator 958_ami_ reverses existing PAH in rats

To assess the downstream molecular functions of NCOA7 activation with 958_ami_, we performed a proximity ligation assay to assess ATP6V1B2-NCOA7 interactions. As expected, application of 958_ami_ significantly induced the number of amplifications per cell, suggesting a molecular enhancement at the level of lysosome (**Figs. 7F−7G**). Direct measure of lysosomal pH using LysoSensor Green DND-189 confirmed enhanced acidification indicated by increased fluorescence intensity (**Fig. 7H**). The maintenance of lysosomal acidification with 958_ami_ similarly prevented induction of *CH25H* under IL-1β, which further corresponded to decreased EC immunoactivation as noted by VCAM1 expression and immune cell adhesion to a monolayer (**Figs. 7I−7M and fig. S10A**).

Next, we sought to confirm if 958_ami_ would protect against endothelial immunoactivation in the proinflammatory monocrotaline PAH rat model. Rats were injected intraperitoneally with DMSO or 958_ami_ (7.5 mg/kg) for 10 days post-monocrotaline loading (**Fig. 7N**). Rodents treated with 958_ami_ had appreciable hepatic or renal toxicity nor alterations in left ventricular function as compared to vehicle controls (**figs. S10B−S10K**). Rats treated with the NCOA7 activator 958_ami_ exhibited decreased CH25H expression with a corresponding attenuation in endothelial VCAM1 expression and CD11b^+^ monocyte infiltration at the pulmonary vessel (**Figs. 7O−7R**). As a result, pulmonary vessels demonstrated decreased muscularization, which corresponded to a significant reduction in both right ventricular hypertrophy and RVSP (**Figs. 7S−7U**). Overall, these data identified 958_ami_ as a novel therapeutic agent that reverses PAH pathogenesis with potential implications related to diseases of immune dysregulation.

## Discussion

By harnessing large-scale, multidimensional genomic and metabolomic analytics with concomitant mechanistic experimentation, we found that NCOA7 regulates lysosomal activity and EC sterol metabolism to function as a homeostatic brake and prevents oxysterol-induced inflammation, EC dysfunction, and PAH. Most notably, the presence of the G allele at SNP rs11154337 results in enhanced *NCOA7* expression, thereby reducing inflammation in PAH and establishing mechanistic proof of the underlying genetic association between SNP rs11154337 and PAH mortality and the metabolomic association between the oxysterol signature and PAH severity. Ultimately, our work establishes a paradigm that links fundamental lysosomal biology and oxysterol metabolism to EC behavior with broad implications in the development of molecular diagnostics and therapeutics in PAH, as well as other inflammatory vascular disorders. More broadly, our discovery platform leveraged here serves as a roadmap in the illustration of the future potential of mechanistic and translational discoveries in genomic and metabolomic medicine as it relates to historically neglected diseases.

Full application of multi–*omics* datasets and their clinical and molecular associations has yet to be fully realized in the discovery of fundamental and translational biology. This is particularly true for rare and orphan diseases, such as PAH, where limitations in the size of cohorts have prevented robust multidimensional –*omics* overlay. Examples of molecular profiling have primarily focused on single molecules or pathways in the search for associations to clinical parameters of risk or disease severity (*63, 64*). Yet, without avenues to pursue cross-validation or to provide mechanistic proof, these narrow pipelines rely upon population-wide significance of association and thus often ignore signals that are biologically relevant but otherwise compromised by a suboptimal statistical power. In this study and our parallel study (*34*), we offer an example in the harmonization of large, unbiased metabolomics with both experimental validation and targeted genomic analyses to define the importance of a SNP-NCOA7-oxysterol axis to PAH. As such, this work offers a model of how gene, protein, and metabolite disease landscapes can be integrated and studied. The continued collaboration to share resources and datasets across multiple, global patient referral centers, as seen recently in PAH datasets (*63, 65*), will further aid the pursuit of multi–omics profiling. Ultimately, the harmonization of datasets, anonymization of clinical records, and continued open access will be essential for continued advancement of multi–omics discovery to personalized medicine in PAH and beyond.

The identification of NCOA7 as a primary controller of PAH has broad implications for human disease. Previously studied in bacterial (*20*) and viral invasion (*17, 21*), renal acidification (*22*), and neuron physiology (*18*), the role of NCOA7 in immunomodulation described in this study may point to much broader actions of NCOA7 isoforms and related proteins. While recent studies have reported unique activity of the short-length isoform of NCOA7 (*17, 18, 21*), our findings describe an additive or synergistic behavior of both isoforms in modulating oxysterol-mediated EC inflammation (**Fig. 4**). The NCOA7 isoforms carry a Tre2/Bub2/Cdc16 (TBC), lysin motif (LysM), domain catalytic (TLDc) domain (*66*). A recent study has demonstrated that all TLDc-containing proteins physically interact with V-ATPases, thereby defining a new class of V-ATPase regulatory proteins (*19*). Moreover, a homologue of NCOA7—oxidation resistance protein 1 (OXR1)—has been implicated in the immune response (*20*), as an anti-inflammatory factor (*67*) presumably through control of lysosomal function (*68*). Thus, future work should address whether TLDc-containing proteins carry compensatory or synergistic activity with NCOA7 in pulmonary vascular disease across lysosomal function and inflammatory cascades.

At the cellular level, our work highlights a broad lysosomal role in EC function and PAH. Prior clinical observations have suggested a relationship between rare, recessive, loss-of-function lysosomal storage disorders and pulmonary vascular diseases. For example, human mutations in various V-ATPase subunits (*e.g.*, *ATP6V1A* and *ATP6V1E1*) can present with pulmonary arterial stenosis or hypoplasia and right ventricular hypertrophy (*16*). PAH has been seen in mucolipidosis—a disease driven by dysfunctional lysosomal enzyme processing (*13–15*). High pulmonary arterial pressures were reported in patients with Gaucher’s disease—a condition resulting from deficiency of lysosomal β-glucosidase (**Figs. 2F and 2G**) and carrying a known association to World Symposium on Pulmonary Hypertension (WSPH) Group 5 PH (*7, 69*). Moreover, Niemann-Pick disease and Fabry disease manifest with severe pulmonary dysfunction (*70*), which often coexist with WSPH Group 3 PH. Supported by these rare genetic diseases, the association of the homozygous C/C genotype to worsened survival in PAH offers broader and more definitive proof of a causative link between lysosomal dysfunction and PAH. In fact, guided by the C mean allele frequency (∼0.48 to 0.52) of SNP rs1115447 in the global population, approximately a quarter of PAH patients harboring the C/C genotype would be expected to suffer from worsened mortality. Furthermore, based on the principle of synergistic heterozygosity (*71*) previously reported in the context of *BMPR2* mutations in familial PAH (*72*), it remains to be seen whether worsened PAH or other lysosomal storage disorders may manifest to an even greater extent in monoallelic carriers of known familial PAH mutations or lysosomal enzyme mutations if accompanied by the SNP rs1115447 C/C genotype.

Our collective work advances the importance of oxysterol and bile acid metabolism to immunoactivation of the endothelium—a poorly explored principle to date (*32*). Emphasizing the relevance of our work to human disease, prior studies have reported elevated oxysterols in patient serum and lung tissue in PAH (*73–75*), and small metabolomics-based studies have demonstrated the elevation of bile acids in PAH plasma and lung tissue (*28–30, 76*). In that vein and consistent with our collective studies of endothelial oxysterol production (*34*), pulmonary tissue, rather than liver tissue, has been described as a major source of circulating extrahepatic bile acids (*77*). Future studies should explore any putative and synergistic pathobiology of the entire complement of oxysterols and bile acids associated with PAH mortality. This includes 25HC, which is already known as both a proinflammatory and anti-inflammatory molecule in innate and adaptive immunity (*31*) and peripheral vascular disease (*78, 79*). Finally, our findings emphasized the relationship of this bile acid and other oxysterols to metrics of disease severity rather than the risk of PAH. As such, in contrast to a single trigger, oxysterol-induced inflammation may act as a “second-hit” to worsen PAH, thus aligning with nuanced evidence of inflammation and immune dysregulation in this complex disease.

Beyond oxysterols, our study also highlights the potential relevance of upstream, systemic cholesterol metabolism in PAH. Notably, reports of HDL and LDL elevations in PAH have been inconsistent (*80–84*). These discrepancies across small studies could be attributed to the complex biology of *de novo* synthesis, uptake, and cellular processing of cholesterol. Alternatively, a more obvious association of HDL and LDL levels may emerge in PAH populations if segregated by the SNP rs11154337 genotype. Inevitably, future work will be necessary to better understand both the systemic and cellular kinetics of total cholesterol metabolism in PAH.

From a translational perspective, the integration of population-level genomic and metabolomic data with mechanistic experimentation offers a compelling foundation for the development of prognostic biomarkers in PAH. Although risk calculation has been implemented in clinical practice (*85, 86*), issues of sensitivity and specificity still plague prognostics with continued reliance on invasive hemodynamic measurements of disease (*87*). As discussed by Alotaibi *et al.* (*34*), assessment of oxysterol and bile acid plasma signatures offers a non-invasive approach to improve risk calculation and guide clinical management. Independently, or in combination, evaluation of SNP rs11154337 carries potential to estimate prognosis and, potentially, to guide more aggressive, targeted therapy. Given the high C and G allele frequencies of this SNP, such genotyping would be relevant to a large portion of PAH patients. Furthermore, considering the broad effects of NCOA7 on inflammation and EC biology, it is possible that other peripheral vascular diseases, such as atherosclerosis, hypertension, pathogen-mediated endotheliopathies like sepsis, and stroke, may associate with this SNP genotype. More recently, the documented relationship between NCOA7 and SARS-CoV-2 cellular entry (*21*), in conjunction with the observed pathophenotype of EC inflammation (*88–90*), suggests that SNP rs11154337 may also predict risk or severity of disease in acute COVID-19 or post-acute sequelae of COVID-19.

Finally, our findings provide a roadmap toward much-needed, next-generation therapies in PAH and conditions of immune dysregulation. Given the causative link between SNP rs11154337 of NCOA7 and EC inflammation, genotyping patients could foster a precision medicine approach in the recruitment and study of ideal candidates for anti-inflammatory therapies that have had mixed results when not guided by such rationale (*91*). This strategy would have direct relevance in endeavors to develop therapeutic agonists of the TLDc domain of NCOA7 or inhibitors of oxysterol and bile acid production to improve existing PAH. Our work offers one such example in the computational modeling of the molecular pocket of NCOA7 in the development of 958_ami_, where both our *in vitro* and *in vivo* studies reveal its abrogation of oxysterol-mediated endothelial immunoactivation and reversal of disease in the PAH monocrotaline rat. Therapies like 958_ami_ would represent a compelling complement to existing vasodilator therapies and other potential agents in development that mainly target cellular proliferative and survival pathways (*92*). Moreover, beyond PAH, it is possible that NCOA7 therapies could modulate adaptive and innate immunity across other diseases where NCOA7 is enriched, such as autoimmune, neurologic, pulmonary, renal, and reproductive contexts.

In summary, via multi-dimensional analyses of genomic and metabolomic datasets in combination with *in vitro* and *in vivo* mechanistic validation, we defined the fundamental and SNP-dependent role of NCOA7 and its control of lysosomal activity and sterol homeostasis to temper inflammation, EC dysfunction, and PAH. Ultimately, our work establishes a paradigm of inflammation linked to lysosomal biology and oxysterol metabolism with wide implications for molecular diagnostic and therapeutic development not only in PAH but also in a multitude of vascular disorders and diseases of immune dysregulation.

## Acknowledgements

This work was supported by NIH grants F30 HL143879 (to L.D.H.); R01 HL124021, HL122596 (to S.Y.C.); American Heart Association Established Investigator Award 18EIA33900027 (to S.Y.C.); R24 HL105333 and R01 HL160941 (to W.C.N.); the French National Research Agency ANR-18-CE14-0025, ANR-21-CE44-0036 and ANR-20-CE14-0006 (to T.B.); the French National Cancer Institute INCA-PLBIO 21-094 (to T.B.). We thank Annie M. Watson for her efforts in patient recruitment. We thank the STRIDE consortium for access to their genomic data funded by NHLBI R01 HL478946 (to R.L.B.) and the NIH–Pharmacogenomics Research Network–Riken Center for Genomic Medicine Collaborative Studies: RIKEN VI (to M.G.-M.). We thank the University of Pittsburgh HSCRF Genomics Research Core (RRID: SCR_018301; Deborah Hollingshead, William Horne, and Janette Lamb) for transcriptomic analyses. We thank the Center for Biologic Imaging at the University of Pittsburgh (to 1S10OD025041-01 C.M.S.C.) for technical assistance and use of their facilities. We thank the Unified Flow Core at the University of Pittsburgh for use of their instruments. We thank the Center for Organ Recovery & Education, the organ donors, and their families for the tissue samples provided for this work. We thank Adam Handen, M.S., Matthew Mitsche, Ph.D., and Robert Farese, M.D. for their expert consultations.

## Methods

### Animal studies

All animal studies were approved by the Division of Laboratory Animal Resources at the University of Pittsburgh. The *Ncoa7* knockout mouse line (C57BL/6 Ncoa7tm1.1(KOMP)Vlcg) was obtained from the Knockout Mouse Project (KOMP; https://komp.org/) and generated using sperm for rederivation at the Genome Editing, Transgenic, & Virus Core at Magee Women’s Research Institute. Obtained mice were bred in-house to generate homozygous, *Ncoa7* knockout mice. To elicit a model of pulmonary inflammation resulting in severe PH, *Ncoa7* knockout mice were crossbred with C56BL/6 Il6 transgenic (Tg^+^) mice. The *Il6* Tg^+^ mice contain a Clara cell 10-kD promoter (CC10) that drives constitutive expression of IL-6 within the lung (*53*). C57BL/6 mice were used for orotracheal delivery of either PBS or 7HOCA (10 mg/kg) serially injected every 5 days for 4 weeks under chronic hypoxia (10% O_2_). Mice were taken to 15 weeks of age under normal oxygen tension before echocardiography, invasive hemodynamics measurement, and tissue harvesting. A monocrotaline rat model of PAH was utilized with a single injection of monocrotaline (80 mg/kg) at 8 to 9 weeks of age. Rats were then injected intraperitoneally with DMSO or 958_ami_ (7.5 mg/kg) for 10 days post-monocrotaline loading dose before takedown.

### Human subjects

Experimental procedures that involved human tissue, plasma, or invasive and noninvasive hemodynamics were approved by the Institutional Review Board at the University of Pittsburgh. Ethical approval for this study and informed consent were consistent to the standards of the Declaration of Helsinki.

### Statistics

All *in vitro* data represent at least three independent experiments. The number of animals used for a given experimental model was calculated to measure at least a 20% difference between the means of the control and experimental groups with a power of 80% and a standard deviation of 10%. The number of patient samples used for molecular analyses was primarily determined by clinical availability. For normally distributed data, paired data were analyzed with a two-tailed Student’s *t*-test and grouped data were compared with either a one- or two-way analysis of variance with post hoc Tukey analysis to adjust for multiple comparisons. Significance was determined by a *P* value less than 0.05. All data are presented as mean ± standard deviation.

Additional methods can be found in the **Supplementary Material**.

## References

1. M. Back, A. Yurdagul, Jr., I. Tabas, K. Oorni, P. T. Kovanen, Inflammation and its resolution in atherosclerosis: mediators and therapeutic opportunities. Nat Rev Cardiol 16, 389–406 (2019).

2. T. J. Guzik, R. M. Touyz, Oxidative Stress, Inflammation, and Vascular Aging in Hypertension. Hypertension 70, 660–667 (2017).

3. P. J. Kelly, R. Lemmens, G. Tsivgoulis, Inflammation and Stroke Risk: A New Target for Prevention. Stroke 52, 2697–2706 (2021).

4. Y. Hattori, K. Hattori, T. Machida, N. Matsuda, Vascular endotheliitis associated with infections: Its pathogenetic role and therapeutic implication. Biochem Pharmacol 197, 114909 (2022).

5. J. H. Jones, R. D. Minshall, Endothelial Transcytosis in Acute Lung Injury: Emerging Mechanisms and Therapeutic Approaches. Front Physiol 13, 828093 (2022).

6. H. M. Otifi, B. K. Adiga, Endothelial Dysfunction in Covid-19 Infection. Am J Med Sci 363, 281–287 (2022).

7. G. Simonneau et al., Haemodynamic definitions and updated clinical classification of pulmonary hypertension. Eur Respir J 53, (2019).

8. W. M. Kuebler et al., A pro-con debate: current controversies in PAH pathogenesis at the American Thoracic Society International Conference in 2017. Am J Physiol Lung Cell Mol Physiol 315, L502–L516 (2018).

9. C. Nathan, A. Ding, Nonresolving inflammation. Cell 140, 871–882 (2010).

10. M. Cao, X. Luo, K. Wu, X. He, Targeting lysosomes in human disease: from basic research to clinical applications. Signal Transduct Target Ther 6, 379 (2021).

11. H. Chichger, S. Rounds, E. O. Harrington, Endosomes and Autophagy: Regulators of Pulmonary Endothelial Cell Homeostasis in Health and Disease. Antioxid Redox Signal 31, 994–1008 (2019).

12. J. A. Mindell, Lysosomal acidification mechanisms. Annu Rev Physiol 74, 69–86 (2012).

13. M. Alfadhel, W. AlShehhi, H. Alshaalan, M. Al Balwi, W. Eyaid, Mucolipidosis II: first report from Saudi Arabia. Ann Saudi Med 33, 382–386 (2013).

14. M. Ishak, E. V. Zambrano, A. Bazzy-Asaad, A. E. Esquibies, Unusual pulmonary findings in mucolipidosis II. Pediatr Pulmonol 47, 719–721 (2012).

15. S. Recla, A. Hahn, C. Apitz, Pulmonary arterial hypertension associated with impaired lysosomal endothelin-1 degradation. Cardiol Young 25, 773–776 (2015).

16. T. Van Damme et al., Mutations in ATP6V1E1 or ATP6V1A Cause Autosomal-Recessive Cutis Laxa. Am J Hum Genet 100, 216–227 (2017).

17. T. Doyle et al., The interferon-inducible isoform of NCOA7 inhibits endosome-mediated viral entry. Nat Microbiol 3, 1369–1376 (2018).

18. E. Castroflorio et al., The Ncoa7 locus regulates V-ATPase formation and function, neurodevelopment and behaviour. Cell Mol Life Sci 78, 3503–3524 (2021).

19. A. F. Eaton, D. Brown, M. Merkulova, The evolutionary conserved TLDc domain defines a new class of (H(+))V-ATPase interacting proteins. Sci Rep 11, 22654 (2021).

20. Z. Wang, C. D. Berkey, P. I. Watnick, The Drosophila protein mustard tailors the innate immune response activated by the immune deficiency pathway. J Immunol 188, 3993–4000 (2012).

21. H. Khan et al., TMPRSS2 promotes SARS-CoV-2 evasion from NCOA7-mediated restriction. PLoS Pathog 17, e1009820 (2021).

22. M. Merkulova et al., Targeted deletion of the Ncoa7 gene results in incomplete distal renal tubular acidosis in mice. Am J Physiol Renal Physiol 315, F173–F185 (2018).

23. V. Negi et al., Computational repurposing of therapeutic small molecules from cancer to pulmonary hypertension. Sci Adv 7, eabh3794 (2021).

24. D. Saygin et al., Transcriptional profiling of lung cell populations in idiopathic pulmonary arterial hypertension. Pulm Circ 10, (2020).

25. J. P. Luzio, P. R. Pryor, N. A. Bright, Lysosomes: fusion and function. Nat Rev Mol Cell Biol 8, 622–632 (2007).

26. A. C. Racanelli, A. M. K. Choi, M. E. Choi, Autophagy in chronic lung disease. Prog Mol Biol Transl Sci 172, 135–156 (2020).

27. M. E. G. de Araujo, G. Liebscher, M. W. Hess, L. A. Huber, Lysosomal size matters. Traffic 21, 60–75 (2020).

28. P. Chouvarine et al., Trans-right ventricle and transpulmonary metabolite gradients in human pulmonary arterial hypertension. Heart 106, 1332–1341 (2020).

29. Y. D. Zhao et al., De novo synthesize of bile acids in pulmonary arterial hypertension lung. Metabolomics 10, 1169–1175 (2014).

30. R. Bujak et al., New Biochemical Insights into the Mechanisms of Pulmonary Arterial Hypertension in Humans. PLoS One 11, e0160505 (2016).

31. J. G. Cyster, E. V. Dang, A. Reboldi, T. Yi, 25-Hydroxycholesterols in innate and adaptive immunity. Nat Rev Immunol 14, 731–743 (2014).

32. P. Qin, X. Tang, M. M. Elloso, D. C. Harnish, Bile acids induce adhesion molecule expression in endothelial cells through activation of reactive oxygen species, NF-kappaB, and p38. Am J Physiol Heart Circ Physiol 291, H741–747 (2006).

33. Y. Meng, S. Heybrock, D. Neculai, P. Saftig, Cholesterol Handling in Lysosomes and Beyond. Trends Cell Biol 30, 452–466 (2020).

34. M. Alotaibi et al., Pulmonary primary oxysterol and bile acid synthesis as a predictor of outcomes in pulmonary arterial hypertension. bioRxiv, 2024.2001.2020.576474 (2024).

35. M. Rabinovitch, C. Guignabert, M. Humbert, M. R. Nicolls, Inflammation and immunity in the pathogenesis of pulmonary arterial hypertension. Circ Res 115, 165–175 (2014).

36. W. J. Kent et al., The human genome browser at UCSC. Genome Res 12, 996–1006 (2002).

37. B. B. Chu et al., Cholesterol transport through lysosome-peroxisome membrane contacts. Cell 161, 291–306 (2015).

38. H. J. Kwon et al., Structure of N-terminal domain of NPC1 reveals distinct subdomains for binding and transfer of cholesterol. Cell 137, 1213–1224 (2009).

39. E. L. Huttlin, et al., The BioPlex Network: A Systematic Exploration of the Human Interactome. Cell 162, 425–440 (2015).

40. M. Merkulova et al., Mapping the H(+) (V)-ATPase interactome: identification of proteins involved in trafficking, folding, assembly and phosphorylation. Sci Rep 5, 14827 (2015).

41. D. E. Johnson, P. Ostrowski, V. Jaumouille, S. Grinstein, The position of lysosomes within the cell determines their luminal pH. J Cell Biol 212, 677–692 (2016).

42. F. K. Harlan et al., Fluorogenic Substrates for Visualizing Acidic Organelle Enzyme Activities. PLoS One 11, e0156312 (2016).

43. J. Marciniszyn, J. A. Hartsuck, J. Tang, Mode of inhibition of acid proteases by pepstatin. J Biol Chem 251, 7088–7094 (1976).

44. Z. Diwu, C. S. Chen, C. Zhang, D. H. Klaubert, R. P. Haugland, A novel acidotropic pH indicator and its potential application in labeling acidic organelles of live cells. Chem Biol 6, 411–418 (1999).

45. F. M. Platt, B. Boland, A. C. van der Spoel, The cell biology of disease: lysosomal storage disorders: the cellular impact of lysosomal dysfunction. J Cell Biol 199, 723–734 (2012).

46. M. J. Clague, S. Urbe, F. Aniento, J. Gruenberg, Vacuolar ATPase activity is required for endosomal carrier vesicle formation. J Biol Chem 269, 21–24 (1994).

47. J. Luo, H. Yang, B. L. Song, Mechanisms and regulation of cholesterol homeostasis. Nat Rev Mol Cell Biol 21, 225–245 (2020).

48. N. Zelcer, C. Hong, R. Boyadjian, P. Tontonoz, LXR regulates cholesterol uptake through Idol-dependent ubiquitination of the LDL receptor. Science 325, 100–104 (2009).

49. A. Beigneux, A. F. Hofmann, S. G. Young, Human CYP7A1 deficiency: progress and enigmas. J Clin Invest 110, 29–31 (2002).

50. F. Luchetti et al., Endothelial cells, endoplasmic reticulum stress and oxysterols. Redox Biol 13, 581–587 (2017).

51. F. A. Masri et al., Hyperproliferative apoptosis-resistant endothelial cells in idiopathic pulmonary arterial hypertension. Am J Physiol Lung Cell Mol Physiol 293, L548–554 (2007).

52. S. Sakao et al., Initial apoptosis is followed by increased proliferation of apoptosis-resistant endothelial cells. FASEB J 19, 1178–1180 (2005).

53. M. K. Steiner et al., Interleukin-6 overexpression induces pulmonary hypertension. Circ Res 104, 236–244, 228p following 244 (2009).

54. E. Rojano, P. Seoane, J. A. G. Ranea, J. R. Perkins, Regulatory variants: from detection to predicting impact. Brief Bioinform 20, 1639–1654 (2019).

55. D. Yang et al., 3DIV: A 3D-genome Interaction Viewer and database. Nucleic Acids Res 46, D52–D57 (2018).

56. R. L. Benza et al., Endothelin-1 Pathway Polymorphisms and Outcomes in Pulmonary Arterial Hypertension. Am J Respir Crit Care Med 192, 1345–1354 (2015).

57. M. Gu, Efficient Differentiation of Human Pluripotent Stem Cells to Endothelial Cells. Curr Protoc Hum Genet, e64 (2018).

58. A. Bakan, N. Nevins, A. S. Lakdawala, I. Bahar, Druggability Assessment of Allosteric Proteins by Dynamics Simulations in the Presence of Probe Molecules. J Chem Theory Comput 8, 2435–2447 (2012).

59. J. Y. Lee, J. M. Krieger, H. Li, I. Bahar, Pharmmaker: Pharmacophore modeling and hit identification based on druggability simulations. Protein Sci 29, 76–86 (2020).

60. T. Sterling, J. J. Irwin, ZINC 15--Ligand Discovery for Everyone. J Chem Inf Model 55, 2324–2337 (2015).

61. J. Sunseri, D. R. Koes, Pharmit: interactive exploration of chemical space. Nucleic Acids Res 44, W442–448 (2016).

62. Z. Kurkcuoglu et al., Performance of HADDOCK and a simple contact-based protein-ligand binding affinity predictor in the D3R Grand Challenge 2. J Comput Aided Mol Des 32, 175–185 (2018).

63. C. J. Rhodes et al., Genetic determinants of risk in pulmonary arterial hypertension: international genome-wide association studies and meta-analysis. Lancet Respir Med 7, 227–238 (2019).

64. C. J. Rhodes et al., Plasma proteome analysis in patients with pulmonary arterial hypertension: an observational cohort study. Lancet Respir Med 5, 717–726 (2017).

65. A. R. Hemnes et al., PVDOMICS: A Multi-Center Study to Improve Understanding of Pulmonary Vascular Disease Through Phenomics. Circ Res 121, 1136–1139 (2017).

66. M. J. Finelli, L. Sanchez-Pulido, K. X. Liu, K. E. Davies, P. L. Oliver, The Evolutionarily Conserved Tre2/Bub2/Cdc16 (TBC), Lysin Motif (LysM), Domain Catalytic (TLDc) Domain Is Neuroprotective against Oxidative Stress. J Biol Chem 291, 2751–2763 (2016).

67. Y. Li et al., Delivering Oxidation Resistance-1 (OXR1) to Mouse Kidney by Genetic Modified Mesenchymal Stem Cells Exhibited Enhanced Protection against Nephrotoxic Serum Induced Renal Injury and Lupus Nephritis. J Stem Cell Res Ther 4, (2014).

68. J. Wang et al., Loss of Oxidation Resistance 1, OXR1, Is Associated with an Autosomal-Recessive Neurological Disease with Cerebellar Atrophy and Lysosomal Dysfunction. Am J Hum Genet 105, 1237–1253 (2019).

69. D. Elstein et al., Echocardiographic assessment of pulmonary hypertension in Gaucher’s disease. Lancet 351, 1544-1546 (1998).

70. R. Borie, B. Crestani, A. Guyard, O. Lidove, Interstitial lung disease in lysosomal storage disorders. Eur Respir Rev 30, (2021).

71. J. Vockley, P. Rinaldo, M. J. Bennett, D. Matern, G. D. Vladutiu, Synergistic heterozygosity: disease resulting from multiple partial defects in one or more metabolic pathways. Mol Genet Metab 71, 10–18 (2000).

72. J. A. Phillips, 3rd et al., Synergistic heterozygosity for TGFbeta1 SNPs and BMPR2 mutations modulates the age at diagnosis and penetrance of familial pulmonary arterial hypertension. Genet Med 10, 359–365 (2008).

73. S. Sharma et al., Role of oxidized lipids in pulmonary arterial hypertension. Pulm Circ 6, 261–273 (2016).

74. A. Al-Husseini et al., Increased eicosanoid levels in the Sugen/chronic hypoxia model of severe pulmonary hypertension. PLoS One 10, e0120157 (2015).

75. D. J. Ross et al., Proinflammatory high-density lipoprotein results from oxidized lipid mediators in the pathogenesis of both idiopathic and associated types of pulmonary arterial hypertension. Pulm Circ 5, 640–648 (2015).

76. Q. J. Han et al., Effects of ranolazine on right ventricular function, fluid dynamics, and metabolism in patients with precapillary pulmonary hypertension: insights from a longitudinal, randomized, double-blinded, placebo controlled, multicenter study. Front Cardiovasc Med 10, 1118796 (2023).

77. A. Babiker et al., Elimination of cholesterol as cholestenoic acid in human lung by sterol 27-hydroxylase: evidence that most of this steroid in the circulation is of pulmonary origin. J Lipid Res 40, 1417–1425 (1999).

78. Z. J. Ou et al., 25-Hydroxycholesterol impairs endothelial function and vasodilation by uncoupling and inhibiting endothelial nitric oxide synthase. Am J Physiol Endocrinol Metab 311, E781–E790 (2016).

79. F. Wang et al., Interferon regulator factor 1/retinoic inducible gene I (IRF1/RIG-I) axis mediates 25-hydroxycholesterol-induced interleukin-8 production in atherosclerosis. Cardiovasc Res 93, 190–199 (2012).

80. N. Al-Naamani et al., Prognostic Significance of Biomarkers in Pulmonary Arterial Hypertension. Ann Am Thorac Soc 13, 25–30 (2016).

81. K. Jonas et al., Highdensity lipoprotein cholesterol levels and pulmonary artery vasoreactivity in patients with idiopathic pulmonary arterial hypertension. Pol Arch Intern Med 128, 440–446 (2018).

82. G. Kopec et al., Low-density lipoprotein cholesterol and survival in pulmonary arterial hypertension. Sci Rep 7, 41650 (2017).

83. C. M. Larsen et al., Usefulness of High-Density Lipoprotein Cholesterol to Predict Survival in Pulmonary Arterial Hypertension. Am J Cardiol 118, 292–297 (2016).

84. S. Umar et al., Involvement of Low-Density Lipoprotein Receptor in the Pathogenesis of Pulmonary Hypertension. J Am Heart Assoc 9, e012063 (2020).

85. M. Humbert et al., Risk assessment in pulmonary arterial hypertension and chronic thromboembolic pulmonary hypertension. Eur Respir J 53, (2019).

86. M. D. McGoon, D. P. Miller, REVEAL: a contemporary US pulmonary arterial hypertension registry. Eur Respir Rev 21, 8–18 (2012).

87. A. Frost et al., Diagnosis of pulmonary hypertension. Eur Respir J 53, (2019).

88. M. Ackermann et al., Pulmonary Vascular Endothelialitis, Thrombosis, and Angiogenesis in Covid-19. N Engl J Med 383, 120–128 (2020).

89. A. Dupont et al., Vascular Endothelial Damage in the Pathogenesis of Organ Injury in Severe COVID-19. Arterioscler Thromb Vasc Biol 41, 1760–1773 (2021).

90. Z. Varga et al., Endothelial cell infection and endotheliitis in COVID-19. Lancet 395, 1417–1418 (2020).

91. R. T. Zamanian et al., Safety and Efficacy of B-Cell Depletion with Rituximab for the Treatment of Systemic Sclerosis-associated Pulmonary Arterial Hypertension: A Multicenter, Double-Blind, Randomized, Placebo-controlled Trial. Am J Respir Crit Care Med 204, 209–221 (2021).

92. M. M. Hoeper et al., Phase 3 Trial of Sotatercept for Treatment of Pulmonary Arterial Hypertension. N Engl J Med 388, 1478–1490 (2023).

