## Supplementary Material for "Genetic regulation and targeted reversal of lysosomal dysfunction and inflammatory sterol metabolism in pulmonary arterial hypertension"

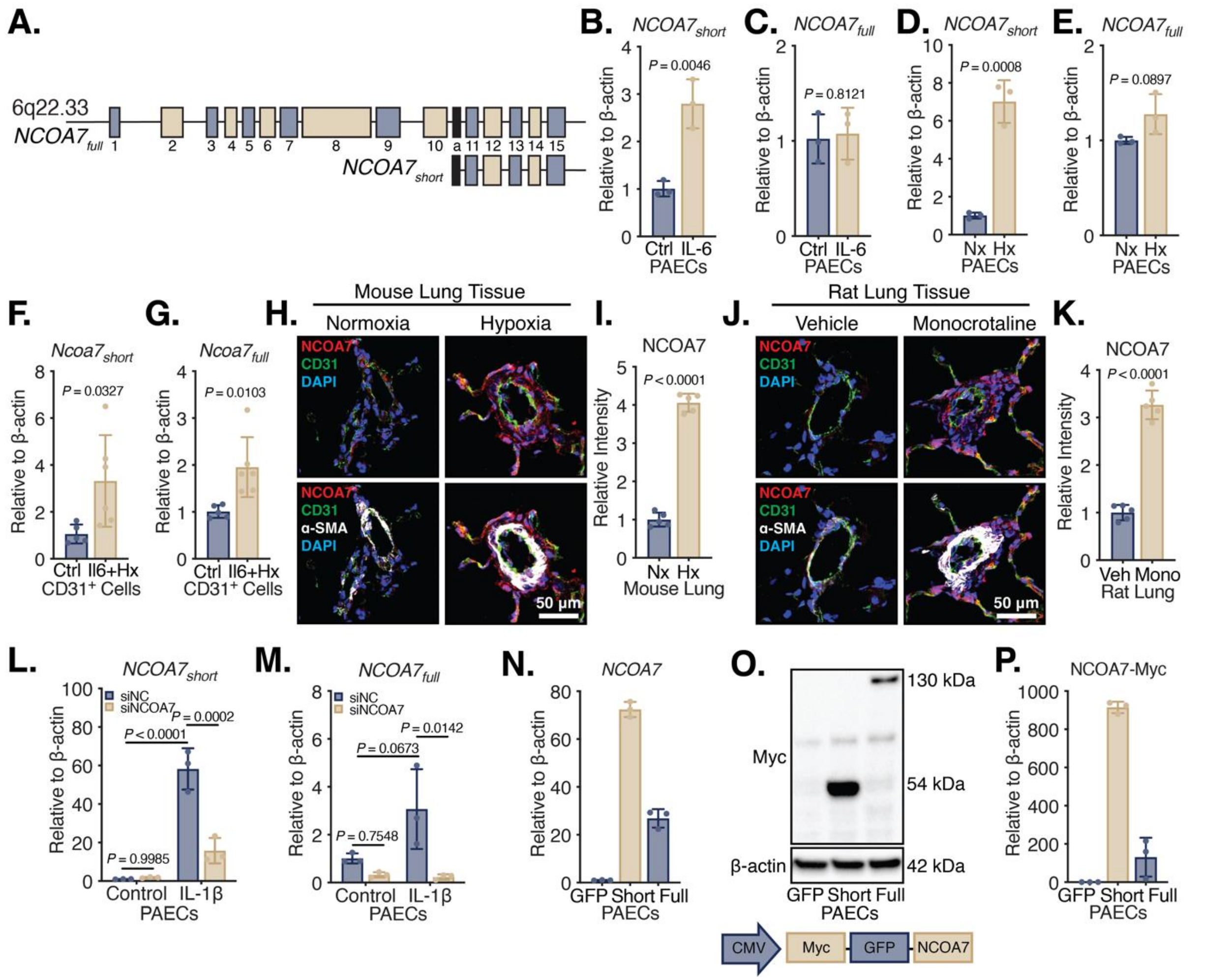


**Fig. S1. Inflammatory regulation of NCOA7 and *in vitro* tools to modulate its expression.** (**A**) Structure of the full-length and short-length isoforms of *NCOA7* located at 6q22.33. Exons are denoted by rectangles. The black rectangle denotes the first exon of the short-length isoform. (**B** to **E**) *NCOA7* isoform expression via RT-qPCR in human PAECs treated with IL-6/IL-6Rα (10 ng/mL) or hypoxia (<1% $O_2$) for 24 hours (N=3/group). (**F** and **G**) *Ncoa7* isoform expression via RT-qPCR in $CD31^+$ isolated cells from total lung of IL-6 transgenic mice under chronic hypoxia. (N=5-7/group). (**H** to **K**) Immunofluorescent staining for and quantification of NCOA7 (red), $CD31^+$ ECs (green), α-$SMA^+$ smooth muscle cells (white), and DAPI-stained nuclei (blue) in pulmonary vessels of mouse (N=5/group) and rat PH models (N=5/group). (**L** and **M**) *NCOA7* isoform expression via RT-qPCR in human PAECs treated with IL-1β and under RNAi against *NCOA7* for

24 hours (1 ng/mL; N=3/group; two-way ANOVA). (**N**) *NCOA7* isoform expression via RT-qPCR human PAECs transduced with a lentivector system delivering either short or full isoform (N=3/group). (**O** and **P**) Immunoblot and densitometry of Myc-tagged NCOA7 isoforms in lentivirus-transduced human PAECs (N=3/group). All data are analyzed by Student's t-test unless otherwise specified and presented as mean ± standard deviation.

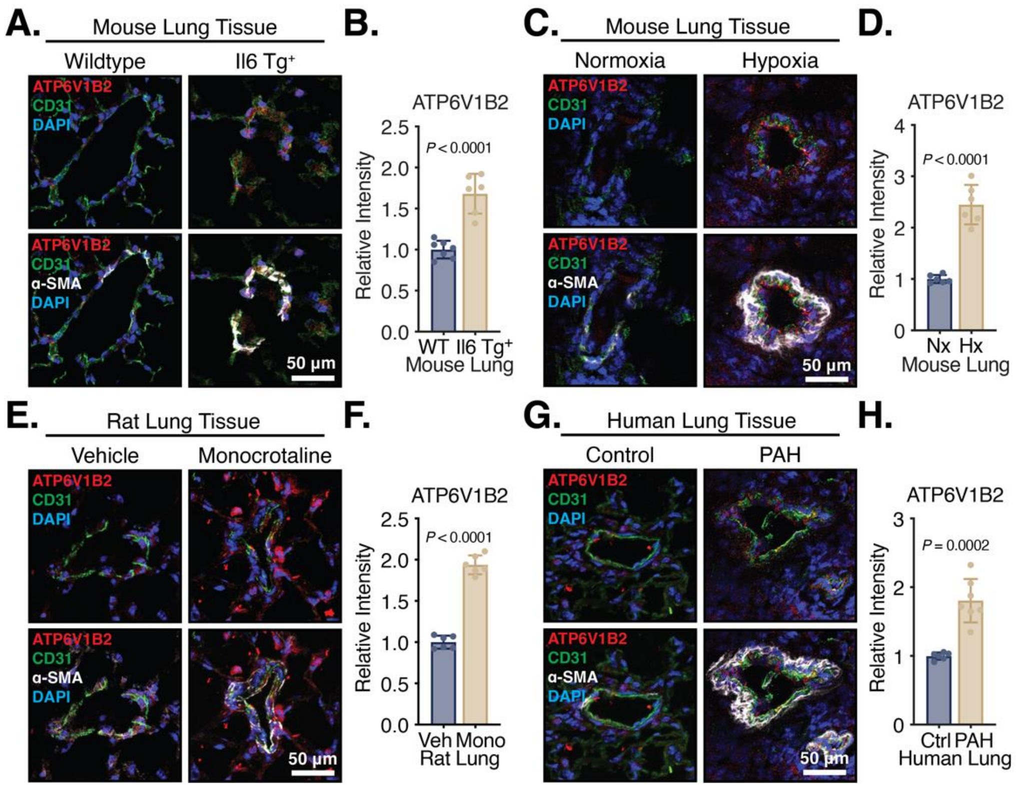


**Fig. S2. NCOA7 binding partner ATP6V1B2 is similarly upregulated in inflammatory models of PH.** (**A** to **H**) Immunofluorescent staining for and quantification of ATP6V1B2 (red), CD31$^+$ ECs (green), α-SMA$^+$ smooth muscle cells (white), and DAPI-stained nuclei (blue) in pulmonary vessels of rodent PH models (N=5-7/group) and humans with PAH (N=6-7/group). All data are analyzed by Student's *t*-test and presented as mean ± standard deviation.

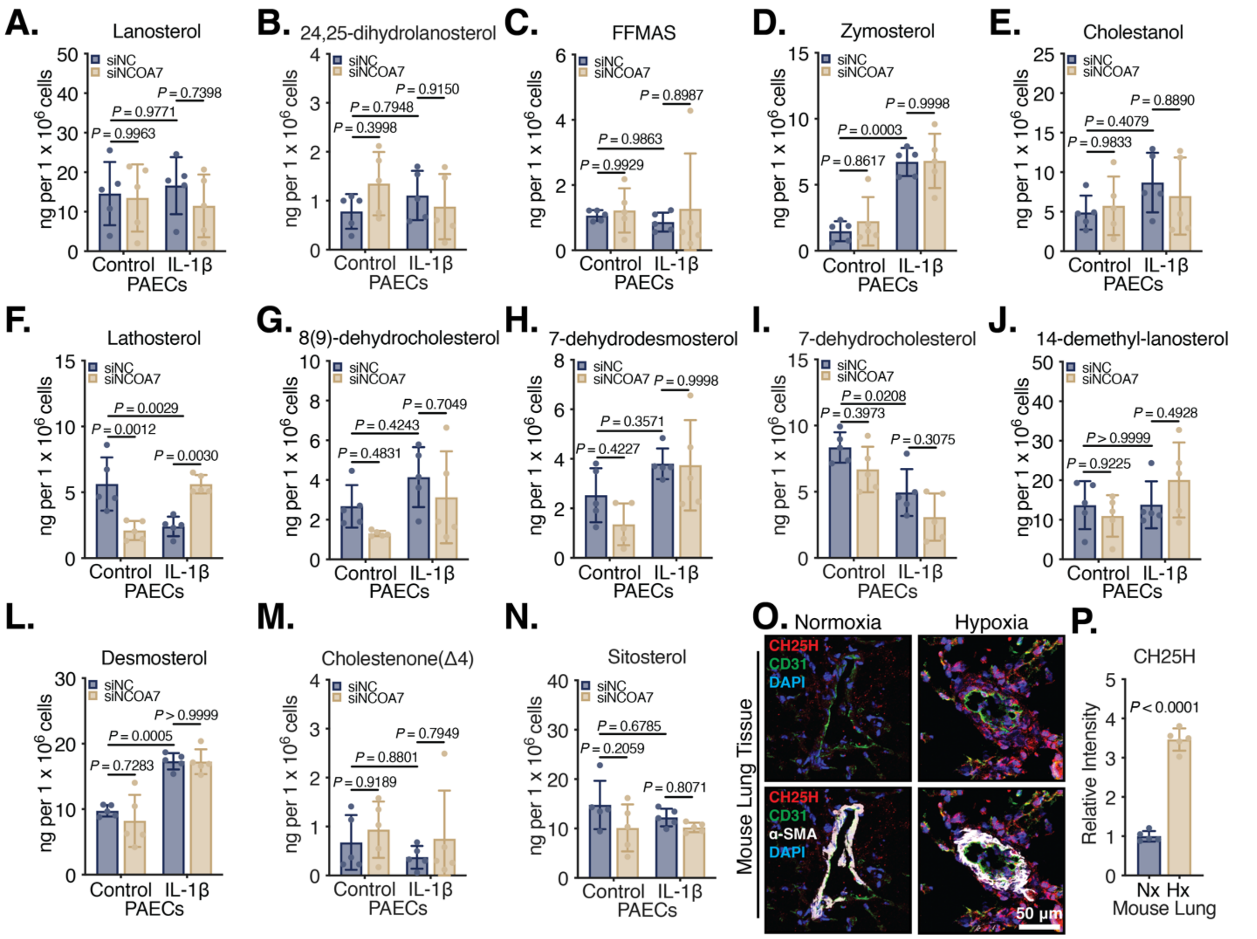


**Fig. S3. NCOA7 modulation of oxysterol and bile acid metabolism is not dependent on sterol flux in the *de novo* synthesis of cholesterol.** (**A** to **N**) Direct measurement of post-squalene intermediates via liquid chromatography-mass spectrometry (N=5/group). (**O** and **P**) Immunofluorescent staining for and quantification of CH25H (red), $CD31^+$ ECs (green), $\alpha\text{-SMA}^+$ smooth muscle cells (white), and DAPI-stained nuclei (blue) in pulmonary vessels of mouse (N=5/group; Student's *t*-test.). All data are analyzed by two-way ANOVA unless otherwise specified and presented as mean ± standard deviation.

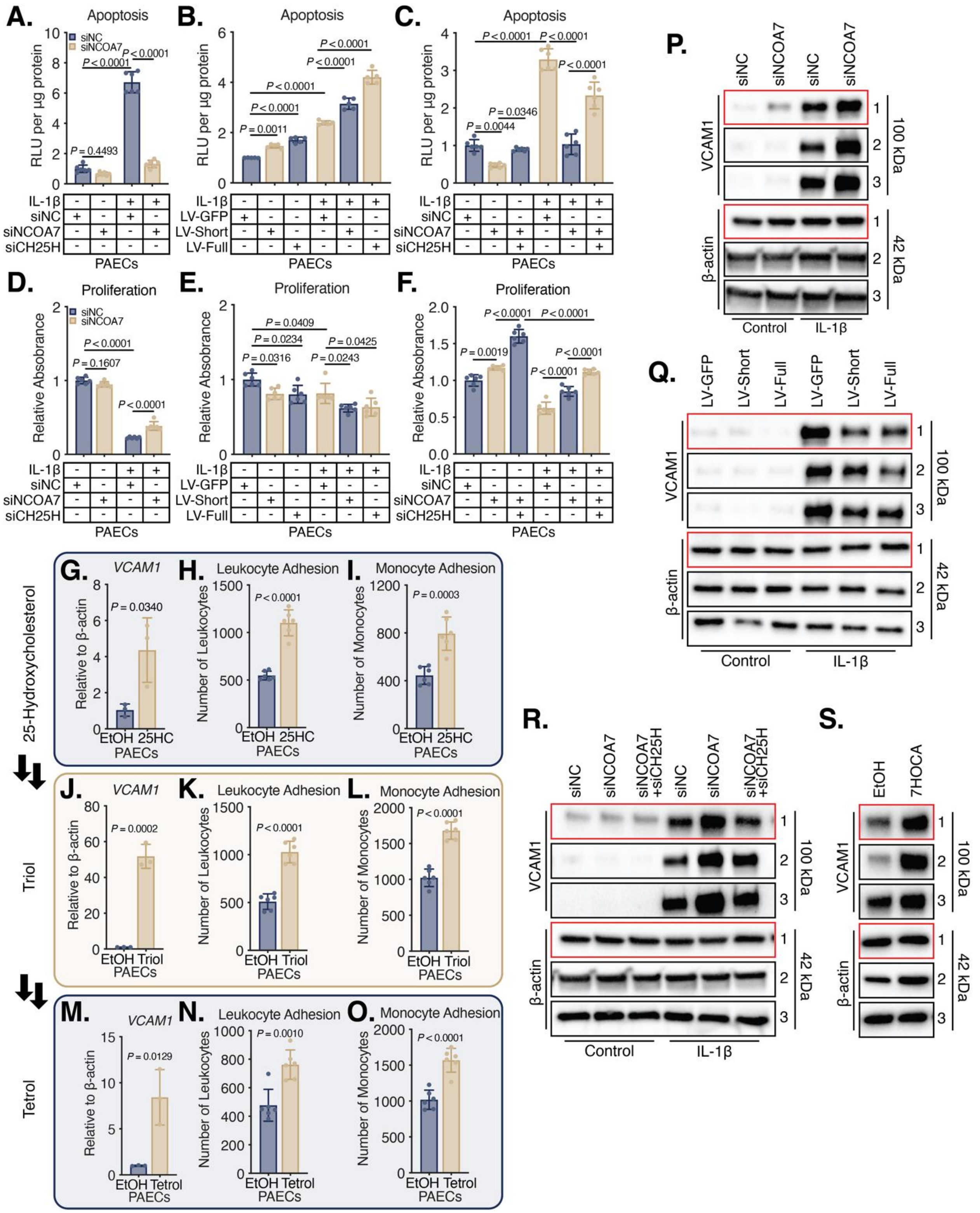


**Fig. S4. NCOA7 modulates endothelial cell apoptosis and proliferation and oxysterols immunoactivated the endothelium.** (**A** to **C**) Apoptosis via caspase-3/7 activity (N=6/group;

Two-way ANOVA). (**D** to **F**) Proliferation via BrdU incorporation (N=6/group; Two-way ANOVA). (**G** to **O**) *VCAM1* expression via RT-qPCR (N=3/group) and leukocyte or monocyte adhesion to a monolayer treated with ethanol versus 25HC (25 μM) or triol (5 μM) or tetrol (50 μM) for 24 hours (N=6/group). (**P** to **S**) Immunoblot of VCAM1 in PAECs in triplicate under various conditions (red box denotes presentation in main figures). All data are analyzed by Student's *t*-test unless otherwise specified and presented as mean ± standard deviation.

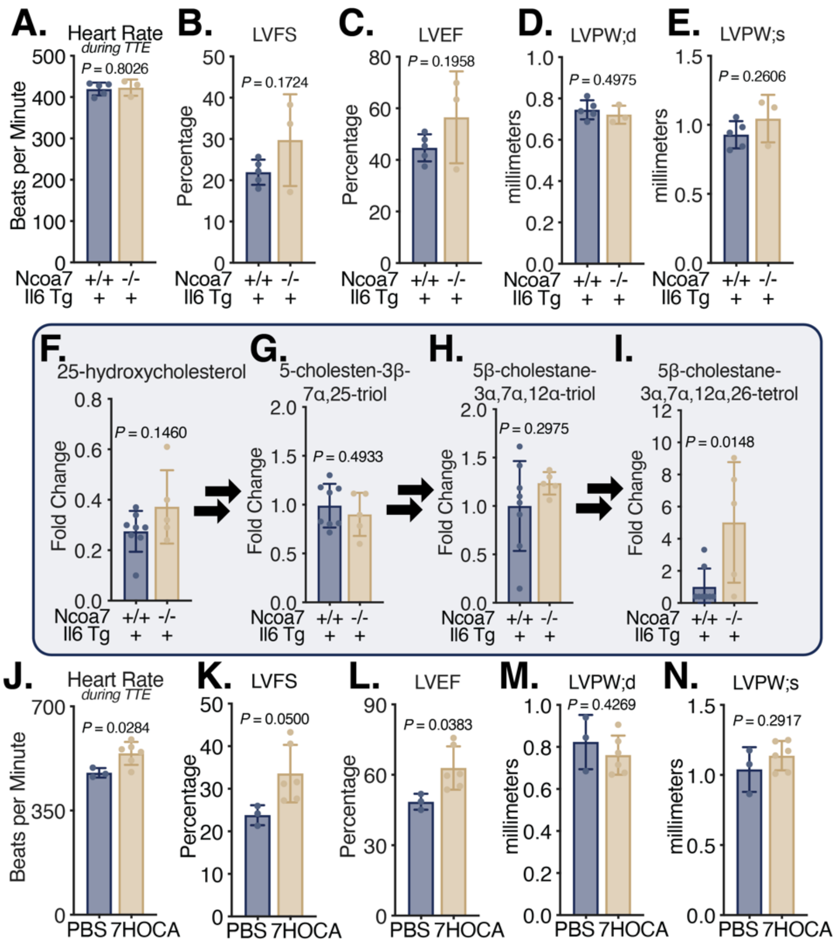


**Fig. S5. Genetic loss of *Ncoa7* does not alter left ventricular function and upregulates plasma oxysterols and bile acids.** Echocardiographic measurements of heart rate, left ventricular fractional shortening (LVFS), left ventricular ejection fraction (LVEF), and left ventricular posterior wall distance during diastole and systole (LVPW;d and LVPW;s) in (**A** to **E**) *Ncoa7*$^{-/-}$ x *Il6* Tg$^{+}$ mice (N=3-5/group) and (**J** to **N**) PBS or 7HOCA (10 mg/kg) mice under four weeks of hypoxia (N=3-6/group). (**F** to **I**) Measurement of oxysterols and bile acids using LC-MS in the serum of *Il6* Tg$^{+}$ versus *Ncoa7*$^{-/-}$ x *Il6* Tg$^{+}$ mice (N=5-8/group). All data are analyzed by Student's *t*-test unless otherwise specified and presented as mean ± standard deviation.

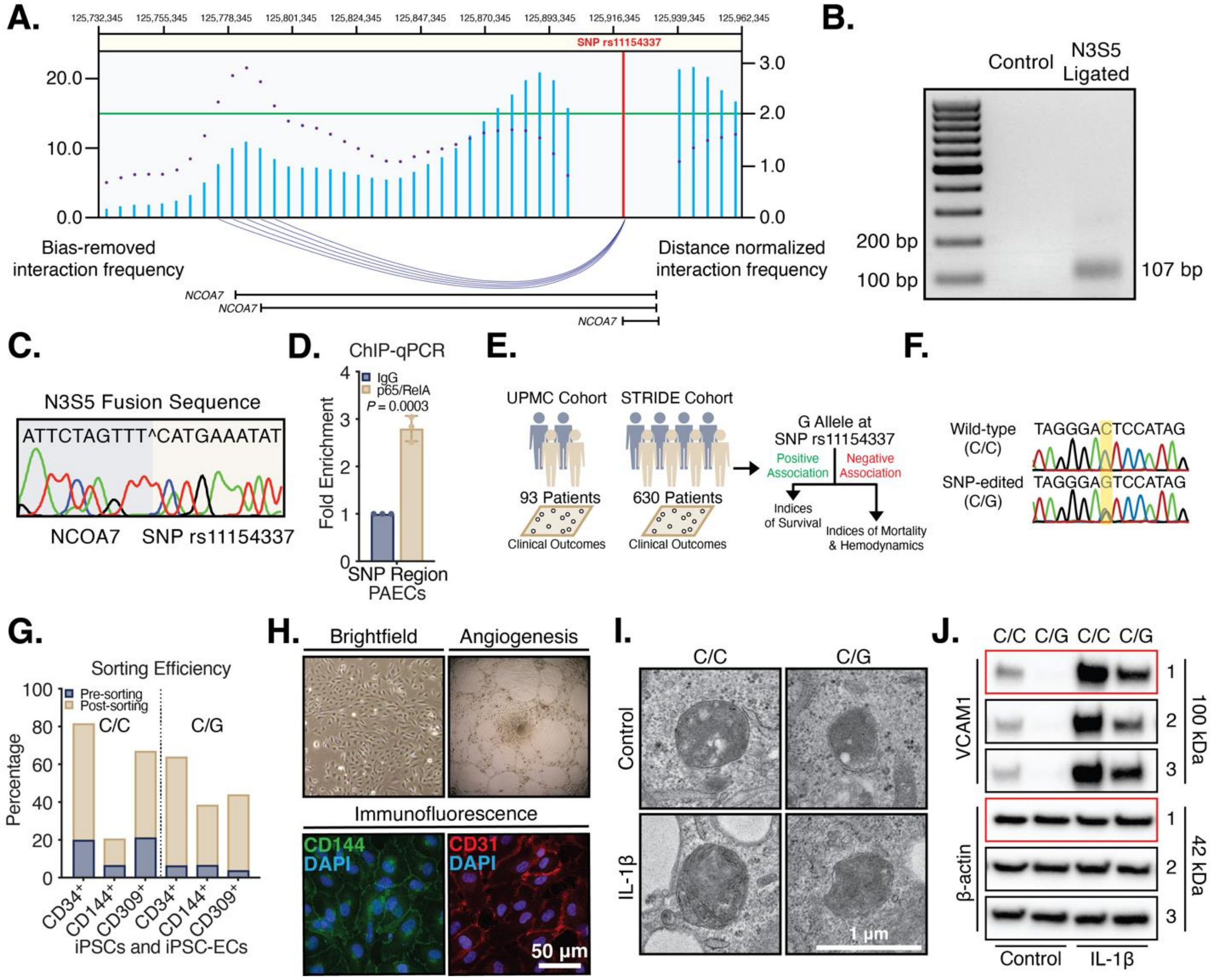

**Fig. S6. Genomic architecture of NCOA7 and the creation of SNP-edited iPSC-derived ECs.** (**A**) High-throughput chromatin conformation capture on human umbilical vein endothelial cells (GEO IDs GSM3438650 and GSM3438651). Blue bars represent the bias-removed chromatin interaction frequency, and the purple dots represent the distance-normalized interaction frequency. The arcs, depicted in indigo, represent the identified interactions of SNP rs11154337 in red defined by the threshold line in green. Data were obtained from the 3D-genome Interaction Viewer & database (3DIV; http://3div.kr). Gene map of NCOA7 isoforms depicted below. (**B**) 3C assay in human PAECs predicting an interaction between the 3' end of restriction enzyme (BspHI) digested genomic DNA fragment containing the promoter of NCOA7 (N3) and the 5' end of genomic DNA fragment containing SNP rs11154337 (S5) that produces a fusion sequence ligated at the BspHI cutting site (N3S5). DNA gel confirming the existence of a 107 bp PCR product with primers targeting the fusion sequence across the BspHI cutting site. The non-ligated genomic DNA was used as control for PCR. (**C**) DNA sequencing of PCR product to confirm the N3S5 fusion sequence. (**D**) ChIP-qPCR of p65/RelA binding to SNP rs11154337 region (N=3/group). (**E**) Schematic of cohorts utilized. (**F**) DNA sequencing of SNP rs11154337 in CRISPR-Cas9, SNP-edited iPSCs. (**G**) Flow cytometry on sorting efficiency of iPSC-derived ECs via MACS. (**H**) iPSC-EC morphology by brightfield, functional capacity by vessel formation, and immunofluorescent staining of EC-specific markers CD144 (green) and CD31 (red). (**I**) Transmission electron microscopy of iPSC-ECs. (**J**) Immunoblot of VCAM1 in iPSC-ECs in triplicate (red box denotes presentation in main figures). All data are analyzed by Student's *t*-test unless otherwise specified and presented as mean ± standard deviation.

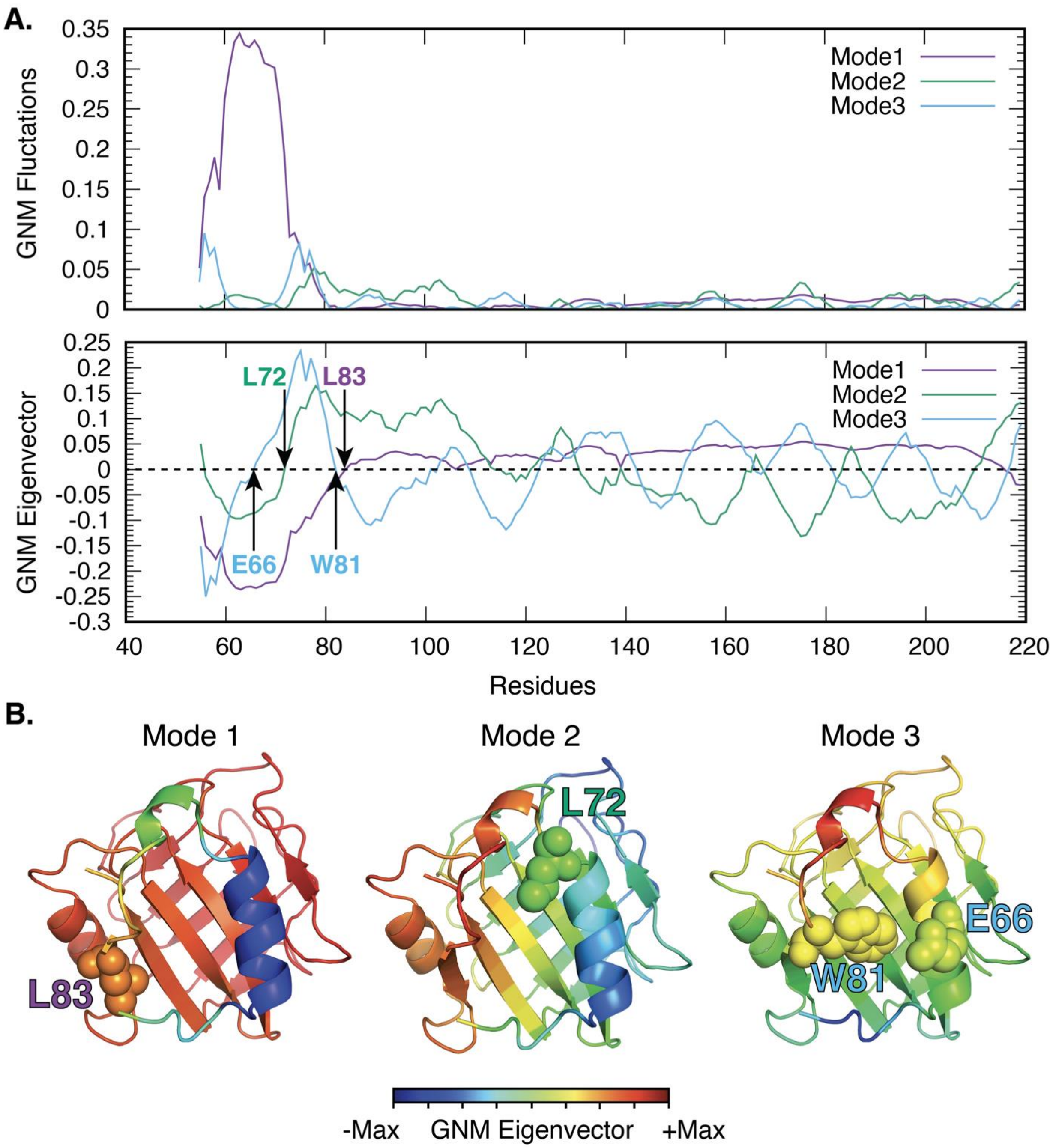


**Fig. S7. Gaussian network model analysis of NCOA7 structural dynamics.** (**A**) Mobility profiles of residues in the most collective three modes of motion of NCOA7 catalytic domain. Dominant hinge residues for each mode are indicated by arrows and labeled. (**B**) Color-coded structures with regions exhibiting largest conformational flexibility are colored in red/blue and minimal flexibility are colored in orange/yellow/green for the three most cooperative modes. Hinge residues for each mode are shown in spheres and labeled.

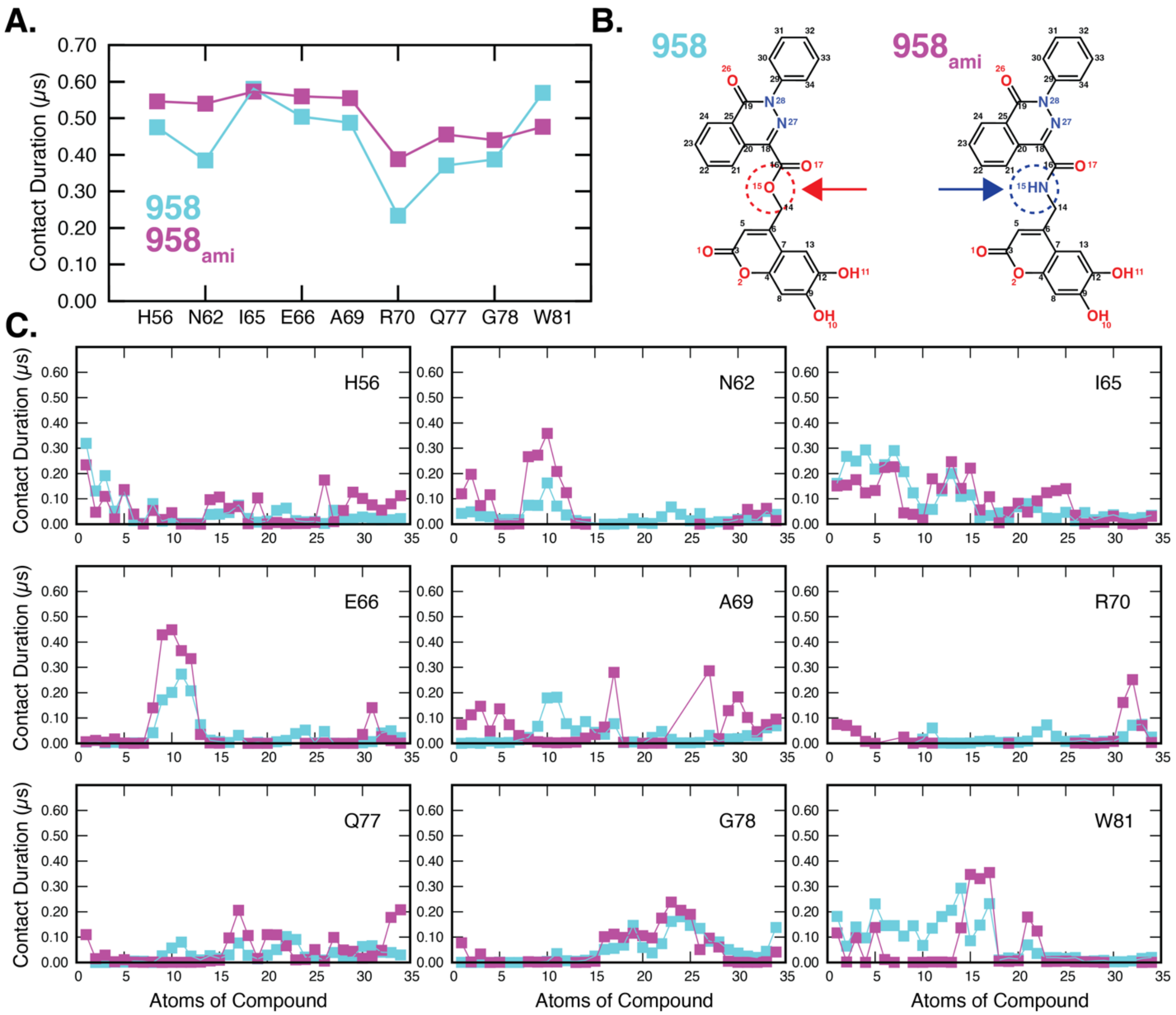


**Fig. S8. Contact duration of the binding of compounds 958 and 958$_{ami}$ to NCOA7 observed in molecular dynamics simulations.** (**A**) Contact duration for NCOA7 residues from MD simulations (three independent runs, each of 0.2 μs; total 0.6 μs for each compound). 958 is shown in *cyan filled-squares* and 958$_{ami}$ in *magenta filled-squares*. Contact is defined as being at a cutoff distance of 4.0 Å between any heavy atoms of the residue and the compound. (**B**) 2D structures of 958 and 958$_{ami}$ with atom ID numbers. (**C**) Nine graphs for contact duration between the specific residues of NCOA7 (listed in X-axis in **A**) and the compound atoms.

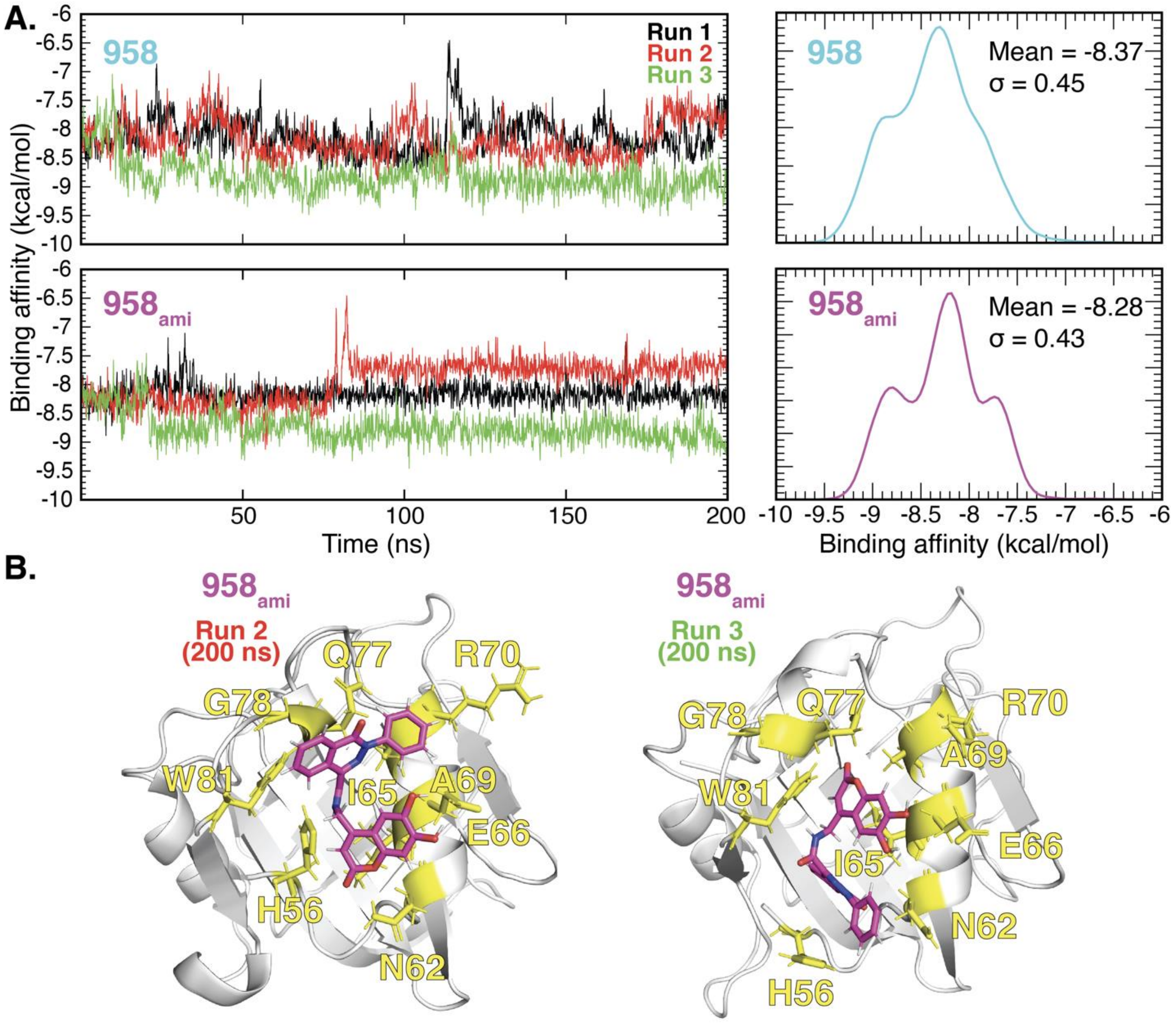


**Fig. S9. Binding affinities observed in MD simulations of NCOA7 complexed with 958 and $958_{ami}$.** (**A**) Time evolution of binding affinities for the two compounds on the *left* panels, referring to three independent runs for each system. The histograms on the *right* are obtained by compiling the snapshots from all three runs for each compound. The average binding affinities (in kcal/mol) and corresponding standard deviations (over the complete duration of the runs) are indicated in each case. (**B**) Binding poses of $958_{ami}$ in respective *run 2* and *run 3* at 200 ns.

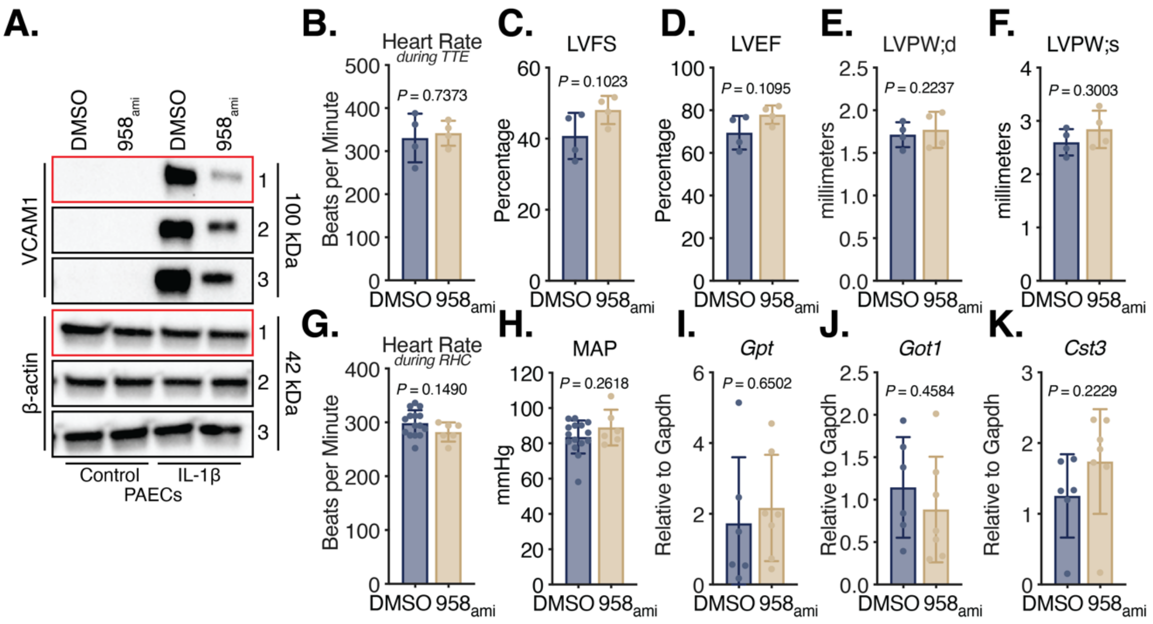


**Fig. S10. $958_{ami}$ does not alter left ventricular function and does not induce hepatic or renal toxicity in rats.** (**A**) Immunoblot of VCAM1 in human PAECs treated with DMSO or $958_{ami}$ in triplicate (red box denotes presentation in main figures). (**B** to **F**) Echocardiographic measurements of heart rate, left ventricular fractional shortening (LVFS), left ventricular ejection fraction (LVEF), and left ventricular posterior wall distance during diastole and systole (LVPW;d and LVPW;s) in monocrotaline rats treated with DMSO or $958_{ami}$ (7.5 mg/kg i.p.) (N=3-4/group). (**G** and **H**) Heart rate and mean arterial pressure (MAP) during right heart catheterization (N=6-15/group). (**I** to **K**) Expression of representative gene transcripts relevant to organ toxicity (*Gpt* and *Got1* in hepatic tissue and *Cst3* in renal tissue) in rats treated with DMSO or $958_{ami}$ (N=5-6/group). All data are analyzed by Student's *t*-test unless otherwise specified and presented as mean ± standard deviation.

| Gene Target | Species | TaqMan Assay ID |
|---|---|---|
| *ACTB* | Human | Hs99999903_m1 |
| *ATP6V1B2* | Human | Hs00156037_m1 |
| *CH25H* | Human | Hs02379634_s1 |
| *LDLR* | Human | Hs01092524_m1 |
| *NCOA7* | Human | Hs00906291_m1 |
| *VCAM1* | Human | Hs01003372_m1 |

**Table S1. TaqMan primers.**

| Gene Target | Species | Primer Sequences 5' to 3' |
|---|---|---|
| *ACTB* | Human | Forward: CATGTACGTTGCTATCCAGGC<br>Reverse: CTCCTTAATGTCACGCACGAT |
| *NCOA7*$_{short}$ | Human | Forward GCCACACTTCTCACTGCTCA<br>Reverse TAGGACAGGCAGCACCTCTT |
| *NCOA7*$_{full}$ | Human | Forward CCCGCTGCAAGATGGAAGG<br>Reverse CGAGTAGCCATCCTGCAACT |
| *Actb* | Mouse | Forward: ACCTTCTACAATGAGCTGCG<br>Reverse: CTGGATGGCTACGTACATGG |
| *Ncoa7*$_{short}$ | Mouse | Forward GCAGGCAACCAGAGAAAGAC<br>Reverse CGTTTTGCCTCCTCAACTGT |
| *Ncoa7*$_{full}$ | Mouse | Forward ATGGAAAGGGTGTGGTTGGG<br>Reverse CTCCAGGCCTGTACAGAGGA |

**Table S2. Custom primers for gene expression.**

| Target | Species | Concentration | Vendor |
|---|---|---|---|
| *Flow Cytometry* | | | |
| CD34-FITC | Mouse | 1:200 | BD Biosciences; 555821 |
| CD144-APC | Mouse | 1:200 | BioLegend; 348508 |
| CD309-PE | Mouse | 1:200 | BD Biosciences; 560872 |
| *Immunoblot* | | | |
| Myc | Rabbit | 1:1000 | Cell Signaling Technologies; 2278S |
| VCAM1 | Rabbit | 1:1000 | abcam; ab134047 |
| *Immunofluorescence* | | | |
| αSMA | Goat | 1:100 | Sigma-Aldrich; A5228 |
| ATP6V1B2 | Rabbit | 1:100 | abcam; ab73404 |
| CD11b | Rabbit | 1:100 | abcam; ab197702 |
| CD31/PECAM-1 | Goat | 1:100 | R&D Systems; AF3628 |
| CH25H | Rabbit | 1:100 | abcam; ab214295 |
| NCOA7 | Rabbit | 1:100 | abcam; ab224481 |
| VCAM1 | Rabbit | 1:100 | abcam; ab134047 |
| *Immunoprecipitation* | | | |
| IgG | Rabbit | 1 μg/μL | abcam; ab171870 |
| NF-κB p65 | Rabbit | 5 μg/μL | abcam; ab19870 |
| *Proximity Ligation Assay* | | | |
| ATP6V1B2 | Rabbit | 1:50 | abcam; ab73404 |
| NCOA7 | Mouse | 1:50 | Santa Cruz Biotechnology; sc-393427 |

**Table S3. Antibodies for protein detection.**

| Oligonucleotides | Species | Sequence 5' to 3' |
|---|---|---|
| *NCOA7* gRNAs | Human | Forward: CACCGTTCAAATATATAGCAGGATA<br>Reverse: AAACTATCCTGCTATATATTTGAAC |
| *NCOA7* ssODN | Human | CTAAACCAAGAAAATGATCATTTGA<br>CAGTGTTTACCTTGGCAAGGATACT<br>GGCCAGGAGGTGTCTTCCCATTAGG<br>GACATGACTATGGACTCACTATCCT<br>GCTATATATTTGAAGACACAGATCAA |
| *NCOA7* Sequencing Primer Set #1 | Human | Forward: CTTCATGGCCTCCTTTGGTA<br>Reverse: AAGGATACTGGCCAGGAGGT |
| *NCOA7* Sequencing Primer Set #2 | Human | Forward: CCAGGTTGAAGTGGAAAGGA<br>Reverse: GGACAGGCCTCACCTGATTA |

**Table S4. Oligonucleotides for genome editing.**

| siRNA Gene Target | Species | ***Silencer*™ Assay ID** |
|---|---|---|
| *CH25H* | Human | 289454 |
| *NCOA7* | Human | 242114 |
| Negative Control No. 1 | Human | AM4611 |

**Table S5. Silencing RNA species.**

| DNA Target | Species | Primer Sequences 5’ to 3’ |
|---|---|---|
| Non-canonical Promoter (*i.e.*, *NCOA7*$_{short}$) | Human | Forward AAAGCTAGGTTCACTGGAGGG<br>Reverse GGCATCGCTGTGAGACTGTAA |
| Canonical Promoter (*i.e.*, *NCOA7*$_{full}$) | Human | Forward TGGGTGGTATGCCTAGTGAA<br>Reverse TTAAGGCTGGGCTGTAAGGT |
| SNP rs11154337 Region | Human | Forward ACCTCCTGGCCAGTATCCTT<br>Reverse ACTCACATAGTGCCCCTCCT |

**Table S6. Custom primers for ChIP qPCR.**

| Software | Access Link |
|---|---|
| EndNote X9.3.3 | https://endnote.com |
| Fiji 2.1.0/1.53c | https://imagej.net |
| FlowJo 10.7.1 | https://flowjo.com |
| LabChart v8.1.14 | https://adinstruments.com |
| Prism 9.1.0 | https://graphpad.com |
| PyMOL 2.3.5 | https://pymol.org/2/ |
| Stata 17.0 | https://www.stata.com |

**Table S7. Software for data analyses.**

### Supplementary Methods

*RNA extraction and quantitative polymerase chain reaction*

Cells were lysed in QIAzol Lysis Reagent (Qiagen; 79306), and RNA was extracted using the Rnaeasy Kit (Qiagen; 74004). Complementary DNA was synthesized using the High-Capacity cDNA Reverse Transcription Kit (ThermoFisher; 4368813). Quantitative real-time PCR (RT-qPCR) was performed on an Applied Biosystems QuantStudio 6 Flex Real-Time PCR instrument. Target gene expression was normalized to a housekeeping gene (*i.e.*, *ACTB*) and fold change was calculated using the $2^{-\Delta\Delta Ct}$ method. TaqMan™ Universal PCR Master Mix (ThermoFisher; 4305719) was used with TaqMan primers (table S1). PowerUp™ SYBR™ Green Master Mix (ThermoFisher; A25742) was used with custom designed primers (table S2).

*Immunofluorescent staining of lung tissue*

OCT-embedded lung tissue was sliced using a cryostat at thickness of 5 to 7 microns and were subsequently mounted onto gelatin-coated histological slides. Sections were rehydrated with PBS for five minutes, fixed in 2% PFA for 30 minutes, permeabilized in 0.1% Triton X-100 for 15 minutes, and blocked in 5% donkey serum and 2% BSA in PBS for one hour at room temperature. Primary antibodies (table S3) were diluted in 2% BSA and incubated overnight at 4°C. AlexaFluor conjugated secondary antibodies were used (ThermoFisher) at a dilution of 1:1000 in 2% BSA for one hour at room temperature. Sections were then counterstained with Hoescht for one minute at room temperature and then mounted. Small pulmonary vessels (30 to 100 microns in diameter) not associated with a bronchial airway were selected for imaging.

*Transfection of cells for RNA silencing*

Human PAECs (Lonza; CC-2530) were transfected at approximately 70 to 80% confluency using negative control (siNC) or target gene (siGene) silencing RNA (siRNA) at 20 nM (table S5). Lipofectamine® 2000 (ThermoFisher; 11668019) was mixed with siRNA per manufacturer's protocol. Lipofectamine®:siRNA mixture was incubated with human PAECs in Opti-MEMTM reduced serum media for 4 to 6 hours (ThermoFisher; 31985062). After incubation, transfection media was removed and replaced with full serum, cell-specific growth media. Experiments were performed 48 hours post-transfection.

*Construction of lentiviral plasmids and particles*

The cDNA sequences encoding full-length NCOA7 (mRNA transcript variant 1, NM_181782.5) and short-length NCOA7 (mRNA transcript 6, NM_001199622.2) were amplified by PCR with HindIII and NheI linkers. The PCR products were directly cloned downstream of Myc-tagged green fluorescent protein (mGFP) open reading frame in the pmGFP-ADAR1-p110 vector (Addgene; 117928). The cDNA sequences encoding mGFP fused full- and short-length NCOA7 were further subcloned into the pCDH-CMV-MCS-EF1α-Puro lentiviral expression vector (Systems Biosciences; CD510B-1). Cloned plasmids were verified by DNA sequencing at the Genomics Research Core at the University of Pittsburgh.

HEK293T cells were maintained in Dulbecco's Modified Eagle Medium (DMEM) with 10% FBS. HEK293T cells were transfected using Lipofectamine® 2000 (ThermoFisher; 11668019) with lentiviral plasmids containing the target gene (or an empty vector for control virus) and packaging plasmids from the ViraPower™ Lentiviral Packaging Mix (ThermoFisher; K497500). Viral particles were harvested 48 hours after transfection, pelleted, and then filtered.

*Transduction of cells for lentiviral vector delivery*

Human PAECs were transduced by direct application of media with polybrene (8 μg/mL) containing viral particles with an empty, control vector expressing GFP or with the target gene. Transduction efficiency was assessed via GFP expression and direct measures of transcript and protein expression. Experiments were performed 72 hours after transduction.

*Protein extraction and immunoblotting*

Cells were rinsed two times with PBS before collection in RIPA buffer containing protease and phosphatase inhibitors. Protein concentration was determined using the PierceTM BCA Protein Assay Kit (ThermoFisher; 23225). Protein lysates were separated using 4–15% Mini-PROTEAN® TGX™ Precast Protein Gels (Bio-Rad Laboratories; 4561086) and subsequently transferred onto a PVDF membrane. Membranes were blocked with 5% BSA in Tris-buffered saline with 0.1% Tween 20 (TBST) for one hour at room temperature. Primary antibodies were subsequently added and incubated at 4°C overnight (table S3). The next day blots were washed three times for 10 minutes each with TBST. Blots were then incubated with the appropriate secondary antibody coupled to HRP for one hour at room temperature. After another round of TBST washing, blots were visualized using Pierce ECL reagents and images were captured using the BioRad ChemiDoc XRS+.

*Chromatin immunoprecipitation and quantitative polymerase chain reaction*

The MAGnify™ Chromatin Immunoprecipitation System (Invitrogen; 49-2024) per the manufacturer’s protocol. Briefly, 1 x $10^6$ human PAECs or iPSC-derived ECs were utilized for each ChIP reaction. Dynabeads® were coupled to either rabbit IgG antibody (1 µg/µL) or rabbit NF-κB p65 antibody (5 µg/µL) for one hour at 4°C (table S3). 1 x $10^6$ cells were trypsinized, pelleted, and resuspended in 500 µL per reaction. Each reaction was crosslinked with 1% methanol-free formaldehyde for 10 minutes at room temperature. The crosslinking reaction was inhibited with 1.25 M glycine for five minutes at room temperature. From this point forward, the reaction was kept at 4°C for all steps. Samples were pelleted and washed three times in cold PBS. Each reaction was then resuspended in 50 µL lysis buffer with protease inhibitor before proceeding to chromatin shearing. The Biorupter® UCD-200 was used to shear cells using a protocol of 20 seconds ON and 40 seconds OFF for six cycles. Samples were pelleted and supernatant containing the chromatin products was collected and confirmed via DNA gel to have appropriate fragmentation. Antibody-bound Dynabeads® were incubated with chromatin for two hours at 4°C while rotating end-over-end. Samples were then washed with a series of immunoprecipitation buffers before crosslinking reversal with proteinase K. The DNA was then purified before proceeding to quantitative PCR. Primers utilized for ChIP-qPCR are listed in table S6.

*Proximity ligation assay*

Direct interaction of NCOA7 with the V-ATPase subunit ATP6V1B2 was assessed using the Duolink® Proximity Ligation Assay (Millipore Sigma; DUO92102). Human PAECs were plated in a Nunc™ Lab-Tek™ II Chamber Slide™ System (20,000 per well; ThermoFisher; 154453) and then fixed with 4% paraformaldehyde for 15 minutes. After permeabilization and blocking per the manufacturer's protocol, wells were incubated with either mouse anti-NCOA7 antibody (Santa Cruz Biotechnology; sc-393427), rabbit anti-ATP6V1B2 antibody (abcam; ab73404), both antibodies, or neither antibodies overnight at 4°C (table S3). Cells were then incubated with the PLUS and MINUS probes for one hour, ligated for 30 minutes, and amplified for 100 minutes at 37°C. Slides were then mounted with ProLong™ Gold Antifade Mountant with DAPI (ThermoFisher; P36935). Images were acquired on a Nikon A1 Confocal Microscope at the Center for Biologic Imaging at the University of Pittsburgh.

*Transmission electron microscopy*

Human PAECs grown on tissue culture plasticware were fixed in 2.5% glutaraldehyde in 100 mM PBS (8 gram/L NaCl, 0.2 gram/L KCl, 1.15 gram/L Na2HPO4.7H2O, 0.2 gram/L KH2PO4, pH 7.4) overnight at 4oC. Monolayers were washed in PBS three times and then post-fixed in aqueous 1% osmium tetroxide, 1% Fe6CN3 for one hour. Cells were washed three times in PBS and then dehydrated through a 30-100% ethanol series with several changes of Poly/Bed® 812 embedding resin (Polysciences). Cultures were embedded by inverting

Poly/Bed® 812-filled BEEM® capsules on top of the cells. Blocks were cured overnight at 37oC, and then cured for two days at 65oC. Monolayers were pulled off the coverslips and re-embedded for cross sectioning. Ultrathin cross sections (60 nm) of the cells were obtained on a Riechert/Leica UltraCut E ultramicrotome, post-stained in 4% uranyl acetate for 10 minutes and then 1% lead citrate for seven minutes. Sections were viewed on a JEOL JEM-1400Flash transmission electron microscope at 80 kV. Images were taken using a bottom mount AMT digital camera. Acquired micrographs were analyzed manually in a blinded manner. Lysosomal area was quantified using Fiji.

*Assessment of lysosomal hydrolase activity*

Lysosomal activity and function were assessed using measures of enzyme activity (*1*). Human PAECs were plated on glass coverslips and stained for all lysosomal measurements. For the LysoLive Assay (Marker Gene Technologies, Inc.; M27745), the β-glucosidase specific substrate GlucGreen was incubated at 5 µM in media for 30 minutes at 37$^{o}$C. Cells were washed three times with ice-cold PBS and subsequently fixed in 4% PFA for 15 minutes at room temperature. Slides were mounted with ProLong™ Gold Antifade Mountant with DAPI (ThermoFisher; P36935).

For the SiR-Lysosome Assay (Cytoskeleton, Inc.; CYSC012), a cell-permeable peptide conjugated to a silicon rhodamine (SiR) dye was incubated in human PAECs as a measure of active cathepsin D. SiR-Lysosome was incubated with cells at 1 µM and with the calcium channel blocker verapamil at 1 µM to improve signal intensity for 30 minutes at 37$^{o}$C. Cells were rinsed three times with ice-cold PBS, fixed in 4% PFA, and mounted as described above.

*Assessment of lysosomal acidification*

Lysosomal acidification was measured using the LysoSensor™ Yellow/Blue DND-160 (PDMPO) dye (ThermoFisher; L7545). The LysoSensor™ Yellow/Blue DND-160 (PDMPO) dye exhibits dual-excitation (i.e., 329 and 384 nm) and dual-emission (i.e., 440 and 540 nm) spectral peaks that are pH-dependent ($pK_a$ 4.2). In acidic organelles, the dye has predominantly yellow fluorescence. In basic organelles, the dye has predominantly blue fluorescence. The unique spectral properties of this dye allow for ratiometric quantification.

Live human PAECs were incubated with 1 µM of dye in 0.1% FBS cell-specific media for 1 minute at 37°C. Cells were rinsed with PBS, trypsinized, pelleted in polystyrene tubes at 300 g for 5 minutes, and rinsed twice more with PBS. Cells were immediately analyzed on a BD LSRFortesssa™ Flow Cytometer (BD Biosciences) at the Unified Flow Core at the University of Pittsburgh. Median fluorescent intensity (MFI) ratio was calculated by using the yellow over the blue MFI fluorescent values.

*Assessment of lysosomal lipid content*

Neutral lipids were stained with the fluorescent dye 4,4-difluoro-1,3,5,7,8-pentamethyl-4-bora-3a,4a-diaza-s-indacene (BODIPY®; ThermoFisher; D3922). Acidic organelles (i.e., lysosomes) were stained with LysoTracker™ Red DND-99 (ThermoFisher; L7528). Live human PAECs were incubated in cell-type specific media containing 1 µM BODIPY® and 50 nM LysoTracker™ Red DND-99 for 30 minutes at 37°C. Cells were then washed with PBS three times before fixation with 4% PFA for 30 minutes at room temperature. Cells were rinsed with PBS three more times and then mounted with ProLong™ Gold Antifade Mountant with DAPI (ThermoFisher; P36935). Images were acquired on a Nikon A1 Confocal Microscope at the Center for Biologic Imaging at the University of Pittsburgh.

Lysosomal lipid content was measured by the degree of colocalization between BODIPY® (i.e., neutral lipids) and LysoTracker™ Red DND-99 (i.e., acidic organelles). Colocalization was measured using EzColocalization in Fiji and quantified as Pearson's Correlation Coefficient (*2*).

*Staining for neutral lipids*

Neutral lipids were stained in cells using the fluorescent dye 4,4-difluoro-1,3,5,7,8-pentamethyl-4-bora-3a,4a-diaza-s-indacene (BODIPY®) as previously described (*3*). Live cells were grown on glass coverslips in plasticware. Cells were rinsed three times with PBS to remove residual media. A staining solution of 2 μM BODIPY® in PBS was applied to cells for 15 minutes at 37°C. Cells were rinsed three times with PBS before fixing with 4% paraformaldehyde for 15 minutes at room temperature. Coverslips were mounted with ProLong™ Gold Antifade Mountant with DAPI (ThermoFisher; P36935). Images were acquired on a Nikon A1 Confocal Microscope at the Center for Biologic Imaging at the University of Pittsburgh.

*Measurement of cholesterol uptake*

To assess cholesterol uptake, a Cholesterol Uptake Assay Kit was utilized per the manufacturer's specifications (abcam; ab236212). Briefly, treated human PAECs were incubated in 0.1% serum, cell-type specific media with supplemented fluorescent, NBD-cholesterol at 20 μg/mL for 24 hours. Cells were rinsed with PBS, trypsinized, pelleted in polystyrene tubes at 300 g for 5 minutes, and rinsed twice more with PBS. Cells were immediately analyzed on a BD LSRFortesssa™ Flow Cytometer (BD Biosciences) at the Unified Flow Core at the University of Pittsburgh. Flow cytometric analysis was chosen over confocal microscopy due to the high rate of photobleaching observed with NBD-cholesterol.

*Assessment of cholesterol content*

To assess cholesterol content, the Cholesterol/Cholesterol Ester-Glo™ Assay Kit was utilized (Promega; J3190). This assay measures cholesterol using a cholesterol dehydrogenase that links the presence of cholesterol to NADH production and thus proluciferin activation. Human PAECs were plated at a density of 20,000 cells per well in a 96-well plate in replicates of six. The assay was performed as the manufacturer specifies.

*Targeted LC-MS for cholesterol intermediates and oxysterols*

Human PAECs were treated and collected for cholesterol intermediates and oxysterol analyses at 1 x $10^6$ cells per glass 16 x 125 mm tube (Pyrex; 9826). Cells were subjected to a liquid-liquid extraction of sterols using a modified form of the Bligh and Dyer method (*4*). Briefly, 1 mL of dichloromethane, methanol, and water were added to each sample. The sample was then vortexed and centrifuged to produce two liquid phases. The lower phase was carefully transferred to a new glass tube and dried under nitrogen. Samples were then resuspended in hexane and the resultant lipid species were analyzed by liquid chromatography-mass spectrometry (LC-MS) at the Center of Human Nutrition at the University of Texas Southwestern Medical Center by Jeffrey McDonald, PhD (*5*). Samples were run on the SCIEX QTRAP 6500+ equipped with a Shimadzu LC-30AD HPLC system and a 150 x 2.1-mm, 5 μm Supelco Ascentis silica column. The LC-MS data was analyzed using MultiQuant (SCIEX).

*Application of oxysterols and bile acids*

Oxysterols and bile acids were applied to human PAECs in 0.1% FBS, cell-specific media. 25-hydroxycholesterol and 7HOCA were dissolved in 100% ethanol and applied to cells at a concentration of 25 μM or 50 μM for 24 hours, respectively.

*Statistical analyses of the UPMC and STRIDE cohorts*

For the UPMC cohort, the effect of SNP rs11154337 G minor allele on six-minute walk distance was calculated using a dominant genetic model. In addition, the effect of the minor allele on time to death or last follow-up was tested using a Cox proportional hazard model. In both models, the genetic effect was adjusted for sex, age, comorbidities, and vasodilator therapies. Stata 17.0 software was used for these analyses.

For the STRIDE cohort, analysis began with raw data in the plink format. Strand information was converted using update_build.sh to InfiniumOmniExpress-24v1-3_A1-

b37.strand. Next, shapeit (*6*) was utilized to phase and impute2 (*7*) with 1000G_Phase3 b37 for imputation. Chromosome bins were merged, and redundant SNPs removed using gtool. First pass of survival analysis was performed with R package gwasurvivr (*8*) performing a coxph test for each SNP versus Time to Death (FinalEvent) including the covariates: sex, age, PAH type, WHO classification, study inclusion (Encysive or Prospective), AnyDrugsBefore, UsePDE, UsePros, UseWarf, UseOxy and a maf filter of 0.005. Further analysis was performed testing SNP rs11154337 against Time to Death (FinalEvent) in patients of European descent (EthConEUR) based on self-reported ethnicity and discriminant principal component analysis.

*Generation of iPSCs and CRISPR-Cas9 gene-editing*

To generate isogenic lines, we introduced the G allele for SNP rs11154337 into control iPSCs with pSpCas9(BB)-2A-GFP (PX458; Addgene; 48138) as the vector for genome editing. For the site-specific CRISPR-Cas9 and guide RNA sequence construction, two reverse complementary guide oligos were annealed and ligated to the linearized PX458 vector (table S4). The single-stranded oligodeoxynucleotide (ssODN) template with mutation/correction sites was designed as previously described (*9*). Control iPSCs were dispersed as single cells the day before transfection at 40 to 50% confluency. The next day, CRISPR-Cas9 and ssODN templates were transfected into iPSCs using GeneJammer reagent according to the manufacturer's protocol (Agilent Technologies 204132). After 36 hours, GFP positive cells were sorted using FACS and were seeded at a density of 2,000 cells per well in a 6-well plate. After expansion for one-week, single cell-derived colonies were observed and picked for genotype characterization and subsequent expansion (table S4). DNA was extracted from collected cells using QuickExtract solution (Epicenter). PrimeSTAR® GXL DNA Polymerase (Clontech) was used for PCR amplification and genotyping of the edited region.

*Differentiation of iPSCs into endothelial cells*

The creation of iPSC-ECs was done using a chemical differentiation protocol with three major steps: mesoderm induction, endothelial specification, and iPSC-EC purification (*10*). Briefly, mesoderm induction was done through Wnt signaling activation using the glycogen synthase kinase-3β inhibitor CHIR99021 (Selleckchem; S2924) in RPMI medium (Life Technologies; 11875-093) with B-27 minus insulin (Life Technologies; A18956-01) supplementation. The use of insulin-free B27 is believed to improve differentiation efficiency, as insulin negatively affects mesoderm induction (*11*). Next, endothelial lineage specification was done using supplemented growth factors like vascular endothelial growth factor (VEGF; 50 ng/mL; Gemini; 300196P) and fibroblast growth factor (FGF; 25 ng/mL; Gemini; 300113P) in EGM™-2 Endothelial Cell Growth Medium-2 BulletKit™. To increase yield, the transforming growth factor β (TGFβ) inhibitor SB431542 (10 μM; Selleckchem; S1067) was also added, as it promotes EC generation and inhibits smooth muscle cell differentiation from endothelial progenitors (*12*). Lastly, iPSC-EC purification was done using magnetic-activated cell sorting (MACS) against the mature EC surface marker vascular endothelial (VE)-cadherin, also known as CD144. Mature iPSC-ECs were labeled with magnetic CD144 MicroBeads (Miltenyi Biotech; 130-097-857), captured by a column in a magnetic field, and then separated from the unlabeled cells. Purified iPSC-Ecs were maintained in EGM™-2 Endothelial Cell Growth Medium-2 BulletKit™.

*Characterization of iPSC-ECs by flow cytometry*

Cells before and after $CD144^+$ purification were analyzed using flow cytometry against endothelial surface markers. Specifically, expression of the mature endothelial progenitor marker CD34 (FITC Anti-CD34 Clone 581; BD Pharmingen™; 555821) (*13*), the mature endothelial marker VE-cadherin/CD144 (APC Anti-CD144 Clone Bv9; BioLegend; 348508) (*14*), and the vascular endothelial growth factor receptor 2 (VEGFR2, or CD309; PE Anti-CD309 Clone 89106; BD Pharmingen™; 560872) (*15*) were assessed (table S3). Samples were run on the BD

LSRFortesssa™ Flow Cytometer (BD Biosciences) at the Unified Flow Core at the University of Pittsburgh.

*Characterization of iPSC-ECs by immunofluorescent staining*

After $CD144^+$ purification, iPSC-ECs were further characterized by immunofluorescent staining of cell surface markers. Briefly, cells were fixed with 4% paraformaldehyde (PFA) for 15 minutes at room temperature, and then blocked in 5% bovine serum albumin (BSA) for one hour at room temperature. Cells were stained with Anti-VE-cadherin/CD144 antibody (1:100; abcam; ab33168) or Anti-CD31 (also known as platelet and endothelial cell adhesion molecule-1; PECAM-1; 1:100; abcam; ab24590) overnight at 4°C (table S3). After rinsing with PBS, cells were incubated with appropriate secondary antibodies (1:1000) in 5% BSA for one hour at room temperature. Cells were rinsed with PBS and mounted with ProLong™ Gold Antifade Mountant with DAPI (ThermoFisher; P36935). Images were acquired on a Nikon A1 Confocal Microscope at the Center for Biologic Imaging at the University of Pittsburgh.

*Characterization of iPSC-ECs by in vitro tube formation*

To confirm an endothelial phenotype, a capillary-like tube formation assay was performed using the in vitro Angiogenesis Assay (R&D Systems; 3470-096-K) (DeCicco-Skinner et al., 2014). A basement membrane extract with reduced growth factors was plated onto a 96-well plate and allowed to solidify for 30 minutes at 37°C. iPSC-ECs were then plated (20,000 cells per well) into the well with cell-type specific media deficient for growth factors and serum. After six hours, capillary-like structures were imaged with the EVOSTM XL Core Imaging System (ThermoFisher) at 10x magnification.

*Chromatin conformation capture (3C) assay*

The 3C assay was performed as previously described (*16*). Briefly, 1 x $10^7$ human PAECs were crosslinked with 1% formaldehyde at room temperature for 10 minutes. Nuclei were isolated and genomic DNA was digested with 400 U BspHI overnight at 37°C, 950 rpm. Per prediction by putative BspHI cutting sites along the genome, the restriction enzyme digestion generates a 1,666 bp genomic DNA fragment containing SNP rs11154337, and a 9,269 bp DNA fragment containing the promoter of NCOA7. Digested DNA was diluted and ligated with or without 4000U T4 DNA ligase for four hours in a 16°C water bath, and then crosslinked DNA was reversed with 100 μg proteinase K at 65°C overnight. DNA was then isolated and purified. PCR was performed with primers chosen to target the potential ligated fusion sequences of the DNA fragment containing SNP rs11154337 and fragment containing the promoter of NCOA7, and close to the BspHI site: (1) 50 bp from the BspHI site at the 3' end of NCOA7 promoter fragment, 5'-TTT GGG CAA TGT TAC AGC AA-3' (forward primer) and (2) 57 bp from BspHI site at the 5' end of the SNP rs11154337 fragment, 5'- GAA ATG CCA GGG ATT CCT TA-3' (reverse primer). The amplified product resulted in a 107 bp fragment to confirm the existence of the fusion sequence. PCR products were separated by gel electrophoresis and analysed by DNA sequencing.

*Leukocyte and monocyte adhesion assays*

Immune activation of the endothelium was assessed by measuring the adhesion of immune cells to an endothelial monolayer. Human PAECs were cultured until a complete monolayer was formed. Immune cells were stained with either CellTrace™ Blue or CFSE (ThermoFisher; Blue, C34568; CFSE, C34554) per the manufacturer's protocol. Between 2.0 to 2.5 x $10^5$ stained immune cells were added to each well of a six-well plate and allowed to incubate for 24 hours. Wells were then rinsed two times with PBS and subsequently fixed with 4% PFA for 15 minutes at room temperature. After fixation, the cells were rinsed once more with PBS. Fluorescent images were acquired at 4x magnification for each well. Immune cell number per image was quantified using Fiji. For the leukocyte adhesion assay, HuT 78 cutaneous T lymphocytes were used (ATCC). For the monocyte adhesion assay, THP-1 peripheral blood monocytes were used (ATCC).

*Apoptosis measured via caspase-3/7 activity*

Apoptosis was assessed using the Caspase -Glo® 3/7 Assay System (Promega; G8090). This assay functions by providing a luminogenic, caspase-3/7 substrate optimized for caspase activity. Cleavage of this substrate generates a luminescence-based signal through luciferase. Equal volumes of this substrate were added to wells containing human PAECs (5,000 per well) and left to incubate at room temperature for 30 minutes. Luminescence was measured via spectrophotometry. Luminescent signal was normalized to protein content per well, assessed using the Pierce™ BCA Protein Assay Kit (ThermoFisher; 23227).

*Proliferation measured via BrdU incorporation*

Proliferation was assessed using the BrdU Cell Proliferation Assay Kit (Cell Signaling Technology; 6813). This assay functions by measuring 5-bromo-2'-deoxyuridine (BrdU) into proliferating cells using an anti-BrdU antibody. BrdU was added to complete growth media containing 5% serum for two hours. Human PAECs (5,000 per well) were fixed and denatured before application of the mouse anti-BrdU antibody. Next, anti-mouse HRP-linked antibody was added. A development substrate was then added to detect with HRP-linked, antibody complexes to BrdU. Absorbance was measured at 450 nm using spectrophotometry.

*Single-cell transcriptomics*

Single cell RNA sequencing was performed on lungs of healthy controls and idiopathic PAH patients. Expression matrices were derived using CellRanger (*17*). Subsequent batch correction, scaling, and normalization were all performed using SCTransform in Seurat v3 (*18-20*). Cell types were determined with SingleR (*21*) using the Blueprint ENCODE reference (*22*).

Cells were identified as positively expressing *NCOA7* if the transformed expression value was greater than 0. Cells expressing *NCOA7* were identified as having a transformed expression value greater than 0.2.

*Transcriptomic analysis of human PAECs and data availability*

Microarray data were obtained using the Affymetrix Clariom D Human Array at the Genomics Research Core at the University of Pittsburgh. The microarray chip was performed on four groups in triplicate for a total of 12 samples. PAECs were subjected to either knockdown control or of the gene *NCOA7*. Additionally, groups were then either left in control conditions or further challenged with the proinflammatory cytokine IL-1β for 24 hours. Raw data were processed using Bioconductor packages in the language R to produce a list of differentially expressed genes that were selected using a Benjamini-Hochberg corrected p-value less than 0.05 in order to minimize the false discovery rate (FDR). The following add-on packages were utilized: oligo, limma, affycore tools, gplots, pd.clariom.d.human for the analysis of this microarray data. Very broadly, the flow of this program serves to take raw microarray data in .CEL format into a list of differentially expressed genes. The workflow included loading the raw data into R, filtering and normalizing the raw data, plotting to demonstrate uniformity among the samples, adding annotations, setting up a comparison matrix to run statistical analyses among the groups, generating a list of differentially expressed genes, filtering said list, and constructing a heat map to depict generalized trends. Gene-set enrichment analyses (GSEA) of identified differentially expressed genes were then created using Gene Ontology (*23, 24*), REACTOME (*25, 26*), KEGG (*27*), and BioCarta (*28*). All microarray data have been submitted to Gene Expression Omnibus (GSE250522).

*Hemodynamic measurements*

Echocardiography was performed on 15-week-old mice using a 15- 45MHz transthoracic transducer and a VisualSonics Vevo770 system (Fujifilm). Anesthesia was administered with 2% isoflurane in 100% $O_2$ during animal positioning and hair removal, and subsequently decreased to 0.8% isoflurane during image acquisition. Data were analyzed in a blinded manner by a technician.

For right heart catheterization, mice were given ketamine/xylazine (9:1; Henry Schein) or subjected to isoflurane (Henry Schein). The isoflurane vaporizer was maintained at 1.5 to 2% with an oxygen gas flow rate of 1 L/min. Right ventricular systolic pressure was measured with Millar catheters (SPR-513 and SPR-671). Catheters were inserted into the jugular vein and then guided through the right atrium and into the right ventricle. Steady right ventricular systolic pressure waveforms were measured for two minutes. Analysis of waveforms were performed in a blinded manner.

*Structural modelling of the catalytic domain of NCOA7$_{short}$*

The sequence of NCOA7$_{short}$ (isoform 5; Q8NI08-5) was downloaded from UniProt (*29*). Its catalytic domain (P55-D219) was modeled using SWISS-MODEL (*30*) and was based on the crystal structure of the TLDc domain of oxidation resistance protein 2 (OXR2) from zebrafish (PDB ID 4ACJ) (*31*). The sequence identity between the catalytic domains of NCOA7 and OXR2 was calculated as 61.8% using Clustal Omega (*32*), which implies that the two domains share the same structure.

*Gaussian network model (GNM) analyses*

In a GNM analysis, the protein structure is represented as an elastic network, where residues serve as nodes of which the positions are identified by those of the α-carbons. As such, the GNM for NCOA7 was developed using a total of 165 residues. The overall potential was represented as the sum of the harmonic potentials between pairs of nodes within an interaction ranged defined as a $C^{\alpha}$-$C^{\alpha}$ distance less than 7.3 Å. The resultant topology of the network was recorded in a N X N Kirchhoff matrix. All computations were performed using the ProDy API (*33, 34*).

*Druggability simulations and analyses*

Druggability simulations were performed for NCOA7 in the presence of the probe molecules, using the all-atom MD simulation package NAMD (*35*) with the CHARMM36 force field for proteins (*36*), the TIP3P water model (*37*), and the CGenFF force field (*38*) for the probe molecules. Probe molecules were benzene, isobutane, imidazole, acetamide, isopropanol, isopropyl amine, and acetate, and were derived from the statistical evaluation of chemical/functional groups most frequently observed in FDA-approved drugs. The trajectories were analyzed using the DruGUI module (*39*) of ProDy (*33, 34*) and, six independent runs of 40 ns were performed. All MD snapshots were superposed onto the reference PDB structure using $C^{\alpha}$-atoms and a cubic grid-based representation of the space with the grid edge size set to 0.5 Å. Probe molecules whose non-hydrogen atoms were within 4.0 Å from protein atoms were considered to interact with the protein. For each probe type, the individual occupancy of grids was calculated using their centroids. We evaluated the occupancy of each probe for each voxel and quantified binding energies using the inverse Boltzmann relation. The resulting binding free energy map identified interaction spots with low (favorable) energy for one or more probe types, and high occupancy voxels were called druggable hot spots.

*Pharmacophore modeling*

Using Pharmmaker, we identified the residues involved in high affinity interactions with each molecular probe type. The residue-probe interactions were subsequently rank-ordered based on their frequency of occurrence at the druggable hot spots in multiple runs (*40*). A snapshot that simultaneously exhibited residue-probe pairings of the probability (*e.g.*, N62-benzene, E66-isopropylamine, P80-benzene, and W81-benzene) was selected as template to construct the pharmacophore model. The pharmacophore model contained a hydrogen bond donor and a hydrophobic feature at the isopropylamine site and hydrophobic and aromatic rings at the two benzene sites. The pharmacophore model was then screened against the ZINC and MolPort libraries using Pharmit (*41*). The MolPort library contains 67,033,884 conformers corresponding to 4,848,718 compounds, and the ZINC library contains 122,276,899 conformers of 13,127,550 compounds (*42*). Among top-scoring compounds, MolPort-004-267-958 was selected for further refinement after initial experimental validation.

*Molecular dynamics simulations of 958 and* $958_{ami}$

All atom MD systems were set-up using GHARMM-GUI Solution Builder (*43*). Simulations were performed using the MD simulation package NAMD (*35*) with the CHARMM36m force field for proteins (*44*), the TIP3P water model (*37*), and the CGenFF force field for the compounds (*38*). Three independent runs of 0.2 μs (for a total 0.6 μs) were performed for 958 and $958_{ami}$. The system in each case comprised the NCOA7 protein and the compound in the presence of explicit water and 0.15 M NaCl, relaxed using the equilibration steps in CHARMM-GUI. We performed NPT dynamics for 0.2 μs with 2 fs time steps, and constant pressure (1 bar) and temperature (300 K). Contact duration between target and compound and hydrogen bond formation were analyzed using VMD 1.9.4 (*45*). Data visualization was performed using PyMOL 2.3.5 and GNUPlot. Binding affinities were calculated using PRODIGY-LIG (*46*).

### References


1. L. V. Albrecht, N. Tejeda-Munoz, E. M. De Robertis, Protocol for Probing Regulated Lysosomal Activity and Function in Living Cells. *STAR Protoc* **1**, 100132 (2020).
2. W. Stauffer, H. Sheng, H. N. Lim, EzColocalization: An ImageJ plugin for visualizing and measuring colocalization in cells and organisms. *Sci Rep* **8**, 15764 (2018).
3. B. Qiu, M. C. Simon, BODIPY 493/503 Staining of Neutral Lipid Droplets for Microscopy and Quantification by Flow Cytometry. *Bio Protoc* **6**,  (2016).
4. E. G. Bligh, W. J. Dyer, A rapid method of total lipid extraction and purification. *Can J Biochem Physiol* **37**, 911-917 (1959).
5. J. G. McDonald, D. D. Smith, A. R. Stiles, D. W. Russell, A comprehensive method for extraction and quantitative analysis of sterols and secosteroids from human plasma. *J Lipid Res* **53**, 1399-1409 (2012).
6. O. Delaneau, J. Marchini, J. F. Zagury, A linear complexity phasing method for thousands of genomes. *Nat Methods* **9**, 179-181 (2011).
7. B. Howie, J. Marchini, M. Stephens, Genotype imputation with thousands of genomes. *G3 (Bethesda)* **1**, 457-470 (2011).
8. A. A. Rizvi *et al.*, gwasurvivr: an R package for genome-wide survival analysis. *Bioinformatics* **35**, 1968-1970 (2019).
9. F. A. Ran *et al.*, Genome engineering using the CRISPR-Cas9 system. *Nat Protoc* **8**, 2281-2308 (2013).
10. M. Gu, Efficient Differentiation of Human Pluripotent Stem Cells to Endothelial Cells. *Curr Protoc Hum Genet*, e64 (2018).
11. X. Lian, J. Zhang, K. Zhu, T. J. Kamp, S. P. Palecek, Insulin inhibits cardiac mesoderm, not mesendoderm, formation during cardiac differentiation of human pluripotent stem cells and modulation of canonical Wnt signaling can rescue this inhibition. *Stem Cells* **31**, 447-457 (2013).
12. D. James *et al.*, Expansion and maintenance of human embryonic stem cell-derived endothelial cells by TGFbeta inhibition is Id1 dependent. *Nat Biotechnol* **28**, 161-166 (2010).
13. C. C. Cheng *et al.*, Distinct angiogenesis roles and surface markers of early and late endothelial progenitor cells revealed by functional group analyses. *BMC Genomics* **14**, 182 (2013).
14. M. Giannotta, M. Trani, E. Dejana, VE-cadherin and endothelial adherens junctions: active guardians of vascular integrity. *Dev Cell* **26**, 441-454 (2013).
15. L. Yang *et al.*, Human cardiovascular progenitor cells develop from a KDR+ embryonic-stem-cell-derived population. *Nature* **453**, 524-528 (2008).
16. E. B. Kabotyanski *et al.*, Lactogenic hormonal induction of long distance interactions between beta-casein gene regulatory elements. *J Biol Chem* **284**, 22815-22824 (2009).
17. G. X. Zheng *et al.*, Massively parallel digital transcriptional profiling of single cells. *Nat Commun* **8**, 14049 (2017).
18. A. Butler, P. Hoffman, P. Smibert, E. Papalexi, R. Satija, Integrating single-cell transcriptomic data across different conditions, technologies, and species. *Nat Biotechnol* **36**, 411-420 (2018).
19. C. Hafemeister, R. Satija, Normalization and variance stabilization of single-cell RNA-seq data using regularized negative binomial regression. *Genome Biol* **20**, 296 (2019).
20. T. Stuart *et al.*, Comprehensive Integration of Single-Cell Data. *Cell* **177**, 1888-1902 e1821 (2019).
21. D. Aran *et al.*, Reference-based analysis of lung single-cell sequencing reveals a transitional profibrotic macrophage. *Nat Immunol* **20**, 163-172 (2019).

22. E. P. Consortium, An integrated encyclopedia of DNA elements in the human genome. *Nature* **489**, 57-74 (2012).
23. C. The Gene Ontology, Expansion of the Gene Ontology knowledgebase and resources. *Nucleic Acids Res* **45**, D331-D338 (2017).
24. M. Ashburner *et al.*, Gene ontology: tool for the unification of biology. The Gene Ontology Consortium. *Nat Genet* **25**, 25-29 (2000).
25. A. Fabregat *et al.*, The Reactome Pathway Knowledgebase. *Nucleic Acids Res*, (2017).
26. D. Croft *et al.*, The Reactome pathway knowledgebase. *Nucleic Acids Res* **42**, D472-477 (2014).
27. M. Kanehisa, M. Furumichi, M. Tanabe, Y. Sato, K. Morishima, KEGG: new perspectives on genomes, pathways, diseases and drugs. *Nucleic Acids Res* **45**, D353-D361 (2017).
28. D. Nishimura, BioCarta. *Biotechnology Software & Internet Journal* **2**, 117-120 (2001).
29. C. The UniProt, UniProt: the universal protein knowledgebase. *Nucleic Acids Res* **45**, D158-D169 (2017).
30. A. Waterhouse *et al.*, SWISS-MODEL: homology modelling of protein structures and complexes. *Nucleic Acids Res* **46**, W296-W303 (2018).
31. M. Blaise *et al.*, Crystal structure of the TLDc domain of oxidation resistance protein 2 from zebrafish. *Proteins* **80**, 1694-1698 (2012).
32. F. Sievers, D. G. Higgins, Clustal Omega for making accurate alignments of many protein sequences. *Protein Sci* **27**, 135-145 (2018).
33. A. Bakan, L. M. Meireles, I. Bahar, ProDy: protein dynamics inferred from theory and experiments. *Bioinformatics* **27**, 1575-1577 (2011).
34. A. Bakan *et al.*, Evol and ProDy for bridging protein sequence evolution and structural dynamics. *Bioinformatics* **30**, 2681-2683 (2014).
35. J. C. Phillips *et al.*, Scalable molecular dynamics with NAMD. *J Comput Chem* **26**, 1781-1802 (2005).
36. R. B. Best *et al.*, Optimization of the additive CHARMM all-atom protein force field targeting improved sampling of the backbone phi, psi and side-chain chi(1) and chi(2) dihedral angles. *J Chem Theory Comput* **8**, 3257-3273 (2012).
37. W. L. Jorgensen, J. Chandrasekhar, J. D. Madura, R. W. Impey, M. L. Klein, Comparison of simple potential functions for simulating liquid water. *The Journal of Chemical Physics* **79**, 926-935 (1983).
38. K. Vanommeslaeghe, A. D. MacKerell, Jr., CHARMM additive and polarizable force fields for biophysics and computer-aided drug design. *Biochim Biophys Acta* **1850**, 861-871 (2015).
39. A. Bakan, N. Nevins, A. S. Lakdawala, I. Bahar, Druggability Assessment of Allosteric Proteins by Dynamics Simulations in the Presence of Probe Molecules. *J Chem Theory Comput* **8**, 2435-2447 (2012).
40. J. Y. Lee, J. M. Krieger, H. Li, I. Bahar, Pharmmaker: Pharmacophore modeling and hit identification based on druggability simulations. *Protein Sci* **29**, 76-86 (2020).
41. J. Sunseri, D. R. Koes, Pharmit: interactive exploration of chemical space. *Nucleic Acids Res* **44**, W442-448 (2016).
42. T. Sterling, J. J. Irwin, ZINC 15--Ligand Discovery for Everyone. *J Chem Inf Model* **55**, 2324-2337 (2015).
43. S. Jo, T. Kim, V. G. Iyer, W. Im, CHARMM-GUI: a web-based graphical user interface for CHARMM. *J Comput Chem* **29**, 1859-1865 (2008).
44. J. Huang *et al.*, CHARMM36m: an improved force field for folded and intrinsically disordered proteins. *Nat Methods* **14**, 71-73 (2017).
45. W. Humphrey, A. Dalke, K. Schulten, VMD: visual molecular dynamics. *J Mol Graph* **14**, 33-38, 27-38 (1996).

46. Z. Kurkcuoglu *et al.*, Performance of HADDOCK and a simple contact-based protein-ligand binding affinity predictor in the D3R Grand Challenge 2. *J Comput Aided Mol Des* **32**, 175-185 (2018).
